# Continuum descriptions of spatial spreading for heterogeneous cell populations: theory and experiment

**DOI:** 10.1101/705434

**Authors:** Oleksii M Matsiaka, Ruth E Baker, Matthew J Simpson

## Abstract

Variability in cell populations is frequently observed in both *in vitro* and *in vivo* settings. Intrinsic differences within populations of cells, such as differences in cell sizes or differences in rates of cell motility, can be present even within a population of cells from the same cell line. We refer to this variability as cell *heterogeneity*. Mathematical models of cell migration, for example, in the context of tumour growth and metastatic invasion, often account for both undirected (random) migration and directed migration that is mediated by cell-to-cell contacts and cell-to-cell adhesion. A key feature of standard models is that they often assume that the population is composed of identical cells with constant properties. This leads to relatively simple single-species *homogeneous* models that neglect the role of heterogeneity. In this work, we use a continuum modelling approach to explore the role of heterogeneity in spatial spreading of cell populations. We employ a three-species heterogeneous model of cell motility that explicitly incorporates different types of experimentally-motivated heterogeneity in cell sizes: (i) monotonically decreasing; (ii) uniform; (iii) non-monotonic; and (iv) monotonically increasing distributions of cell size. Comparing the density profiles generated by the three-species heterogeneous model with density profiles predicted by a more standard single-species homogeneous model reveals that when we are dealing with monotonically decreasing and uniform distributions a simple and computationally efficient single-species homogeneous model can be remarkably accurate in describing the evolution of a heterogeneous cell population. In contrast, we find that the simpler single-species homogeneous model performs relatively poorly when applied to non-monotonic and monotonically increasing distributions of cell sizes. Additional results for heterogeneity in parameters describing both undirected and directed cell migration are also considered, and we find that similar results apply.

## 1 Introduction

*In vitro* cell migration experiments play an important role in the discovery and testing of putative drug treatments, the study of malignant tumour growth and metastasis, as well as tissue regeneration and repair (Savla et al., 2004; Sengers et al., 2007; Tremel et al., 2009; Sarapata and de Pillis, 2010; Gerlee, 2013; Edmondson et al., 2014; Shah et al., 2016). Mathematical models of many biological processes involved in these experiments normally require certain assumptions to make the problem mathematically and computationally tractable. When modelling large populations of cells, one of the most intuitive approaches is to assume that all cells have fixed properties, such as assuming all cells have constant size and constant diffusivity (Sherratt and Murray, 1990; Galle et al., 2005; Simpson et al., 2013). In this framework a cell population is considered to be a *homogeneous* population, and single-species homogeneous models are routinely invoked (Maini et al., 2004a; Maini et al., 2004b; Sepulveda et al., 2013; Simpson et al., 2013; George et al., 2017; Vo et al., 2015). Single-species homogeneous models are much less computationally expensive than more elaborate multi-species heterogeneous models and, as a result, are frequently used relative to multi-species counterparts. In addition, multi-species frameworks usually involve a significantly larger number of free model parameters that we may have little prior knowledge about and so the process of calibrating multi-species heterogeneous models to match experimental observations is significantly more challenging than calibrating single-species homogeneous models. This is an important consideration because it is well-known that parameterising mathematical models of biological processes can be challenging, often requiring computationally-intensive methods (Pozzobon and Perré, 2018; Warne et al. 2019).

Although *heterogeneity* in cell populations is frequently observed in experiments, there is relatively little guidance or consensus in the literature about how to incorporate such heterogeneity into the mathematical models used to replicate and predict such experiments (An et al., 2001; Altschuler et al., 2010; Menon et al., 2018). Figure 1(a)-(b) shows a typical experiment where we can clearly visually observe cells of different sizes. The measured cell size distribution in Figure 1(c) quantifies this heterogeneity in cell sizes and raises the question if the most straightforward approach of applying a single-species homogeneous model can be reasonably used to predict the spatial spreading of this clearly heterogeneous population. In addition to the clear visual heterogeneity in cell sizes, it could be relevant to consider that cells of different sizes can exhibit different behaviour such as different rates of motility, or different mechanical properties including resistance to deformation and adhesion. Therefore, it could be possible that there are multiple types of heterogeneity acting in even this very simple experiment. Previously, heterogeneity in cell populations has been introduced in both discrete and continuum models of cell motility (Simpson et al., 2014; Jin et al., 2016b; Sundstrom et al., 2016; Matsiaka et al., 2017). Previous work has also attempted to estimate parameters in heterogeneous models that describe glioblastoma progression (Rutter et al., 2018). However, these previous modelling studies do not address the basic question of identifying whether it is absolutely necessary to apply a multispecies heterogeneous models to obtain a faithful description of the behaviour of the heterogeneous population and whether different forms of heterogeneity affect the answer to this fundamental question.

**Fig. 1.**
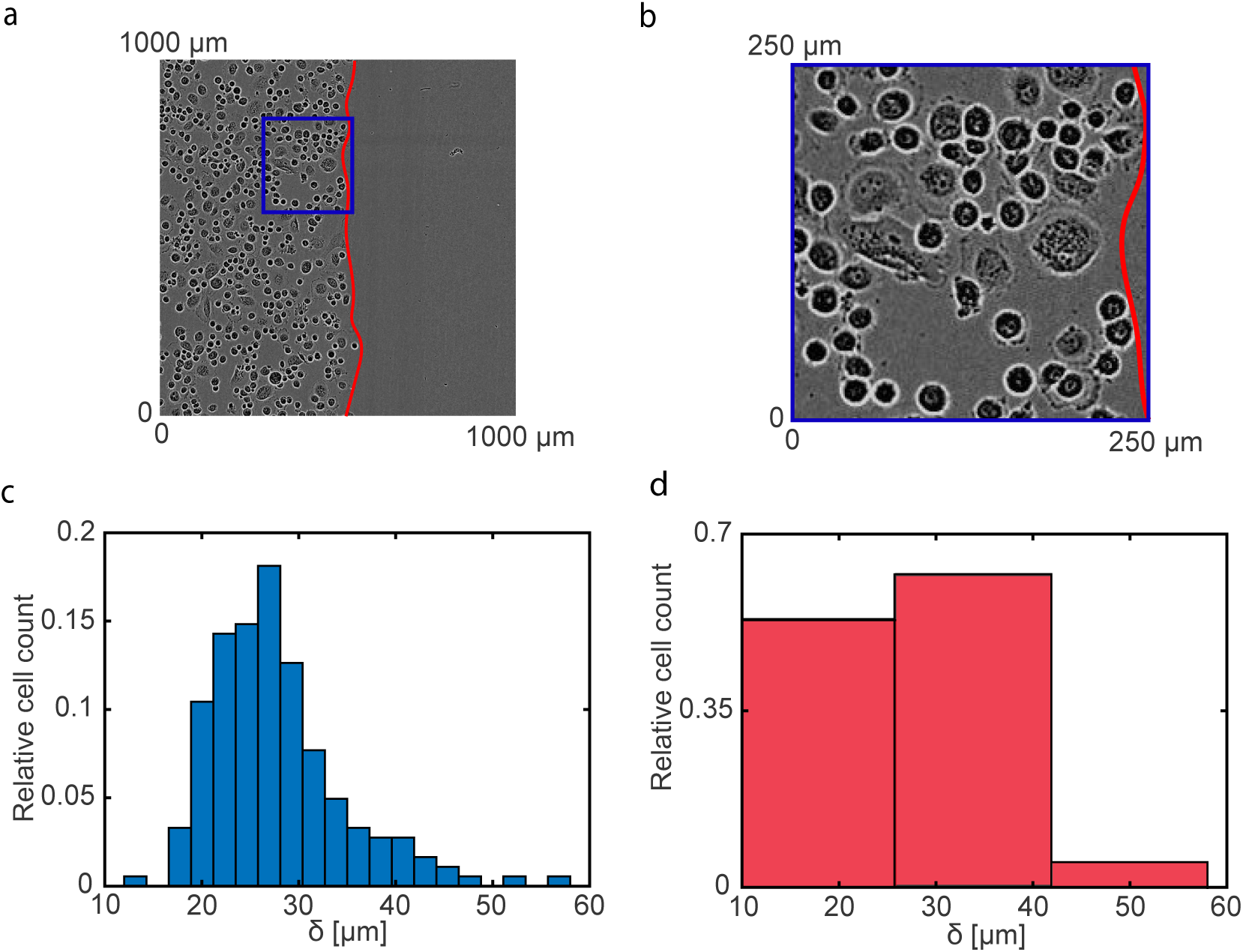
Heterogeneity in a population of PC-3 prostate cancer cells (Kaighn et al., 1979). (a) Experimental image of an advancing cell population and corresponding cell size distribution. The red solid line denotes position of the leading edge. (b) Detailed image of the subregion denoted in the blue rectangle in Figure 1(a). (c) Cell size distribution with a bin size of 15 *µ*m. The cell size distribution is obtained from the sample of 184 cells randomly selected from the population. (d) Cell size distribution with a bin size of 2.3 *µ*m. The histogram in Figure 1(d) is constructed using the same sample of 184 cells.

**Fig. 2.**
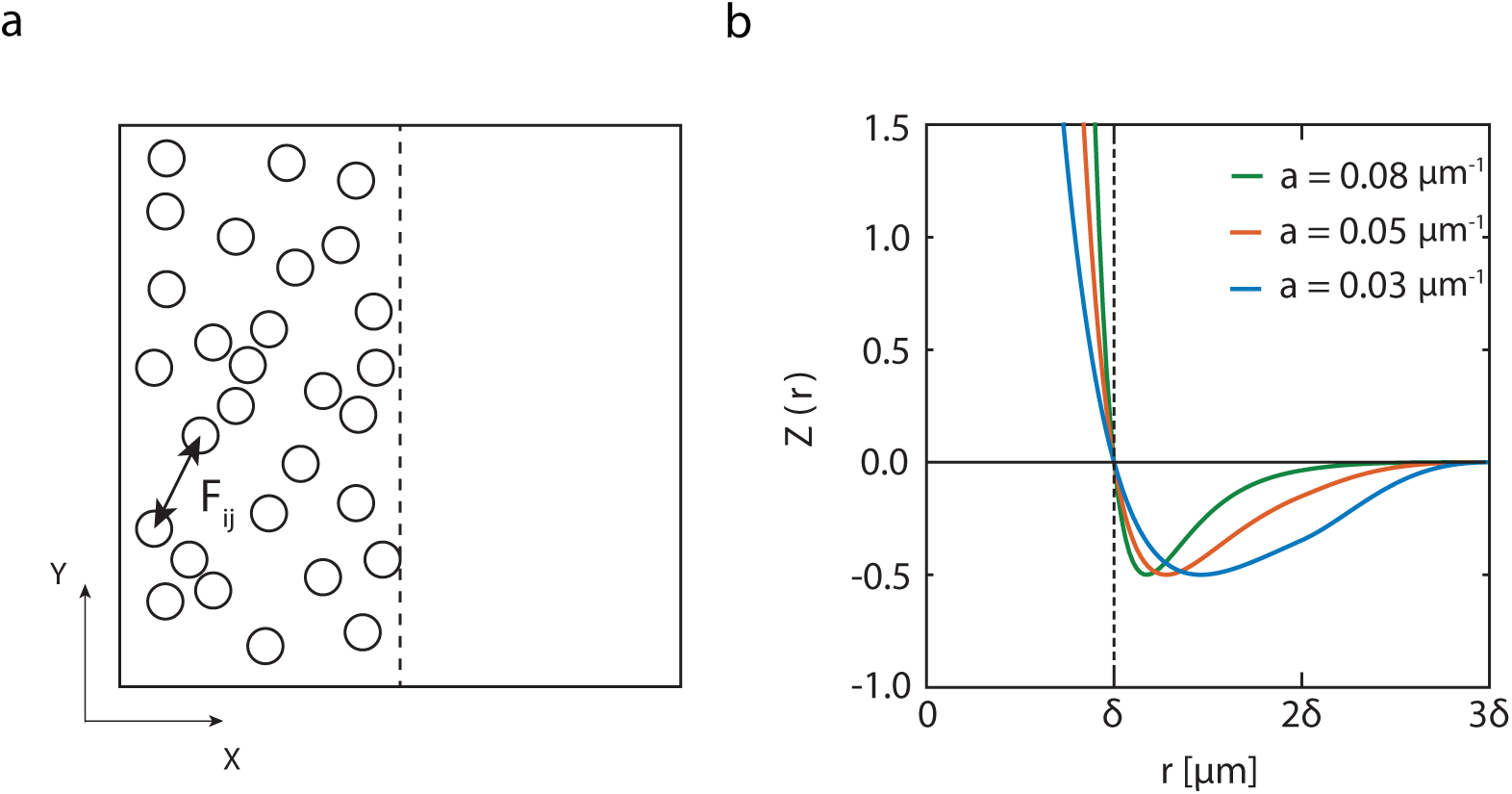
(a) An idealisation of the front-like distribution of cells in the experimental design shown in Figure 1(a). Here all cells are of constant size. *F*_*ij*_ is the interaction force between cell *i* and cell *j*. The vertical dashed line represents the approximate leading edge of the population. (b) A typical cell-to-cell interaction force function in the form of the modified Morse potential, *Z*(*r*), (Equation (3.7)) used to mimic adhesion and repulsion between individual cells. The vertical dashed line represents the diameter of individual agents, *δ*. The horizontal line at *Z*(*r*) = 0 shows the change from long-range attraction (*Z*(*r*) *<* 0 for *r > δ*) to short-range repulsion (*Z*(*r*) *>* 0 for *r < δ*).

In our work we use an experimentally-motivated approach to investigate the role of heterogeneity in two-dimensional scratch assays, and we compare the performance of a single-species homogeneous model relative to a heterogeneous multi-species model. We use numerical solutions of the multi-species heterogeneous model to produce synthetic test data that we use to investigate the performance of a simpler single-species homogeneous model. To mimic experimental data, such as depicted in Figure 1, we use the multi-species continuum approach introduced by Matsiaka et al. (2017). To keep our work tractable, we describe the heterogeneity by dividing the total population into three subpopulations with varying properties. The choice of working with three subpopulations allows us to keep the model computationally tractable while capturing important differences in the population properties, as illustrated in Figure 1(d). Throughout this work we consider four distinct distributions of cell sizes: (i) monotonically decreasing (Set Ia); (ii) uniform (Set Ib); (iii) non-monotonic (Set Ic); and (iv) monotonically increasing (Set Id). The monotonically decreasing distribution, as shown in Figure 3(a), is a fairly accurate approximation of the experimentally observed cell size distribution in Figure 1(d). The other three kinds of distributions are included in our work for completeness. Our findings suggest that, for certain cell size distributions, namely monotonically decreasing and uniform distributions, the single-species homogeneous model performs remarkably well with an excellent match between the density profiles generated by the three-species heterogeneous model and density profiles predicted by its single-species homogeneous analogue. Therefore, our results imply that applying a single-species homogeneous model to describe experiments with monotonically decreasing or uniform cell size distributions might be sufficient for accurately predicting population-level behaviour. In contrast, the data with non-monotonic and monotonically increasing cell size distributions might require the application of multi-species models to account for differences in population.

**Fig. 3.**
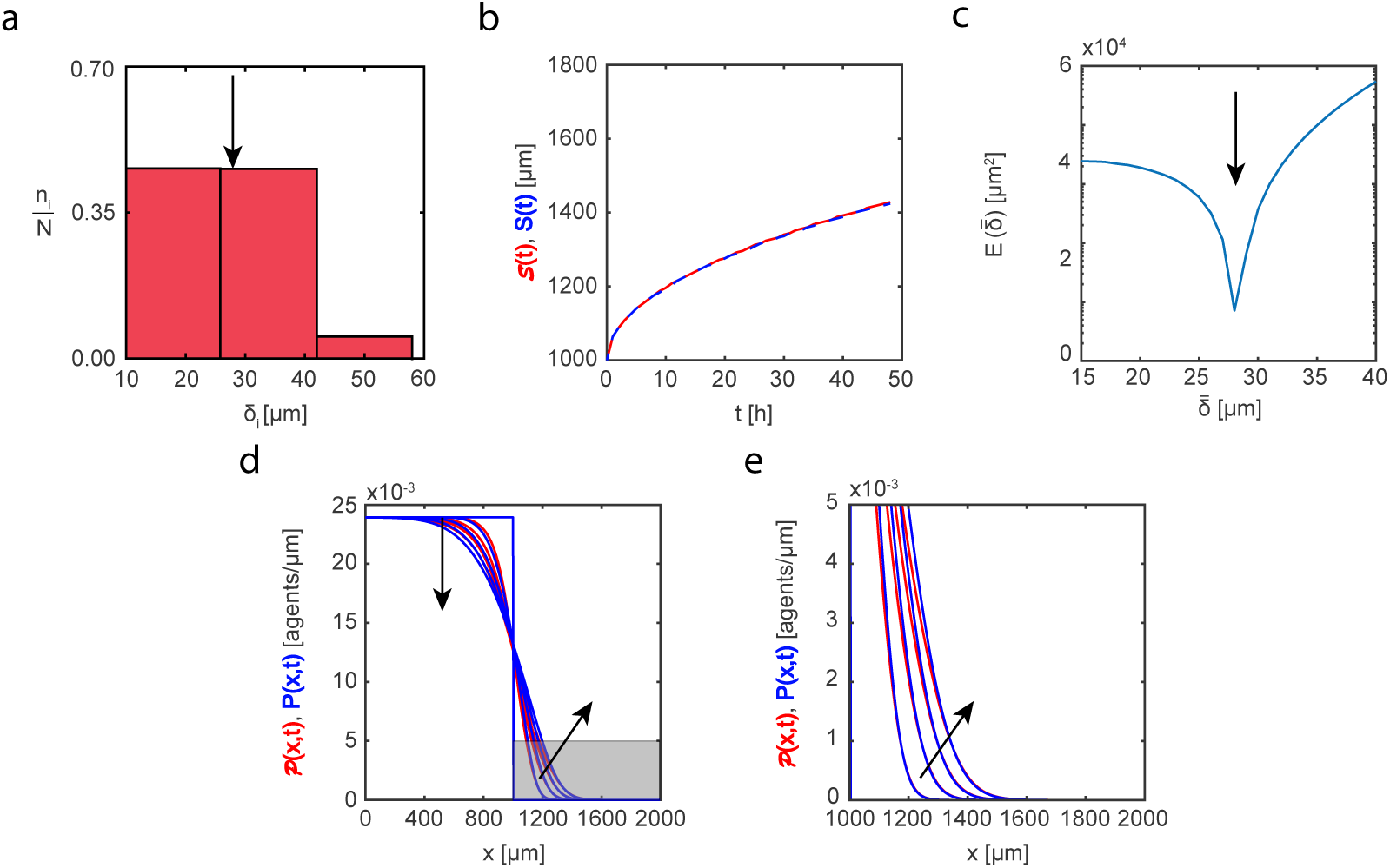
Set Ia. Heterogeneity in cell sizes: monotonically decreasing distribution. (a) Cell size distribution adopted in the three-species heterogeneous model, Equations (3.2)-(3.4). Here the proportions of cells of different sizes are set to: (i) *n*_1_*/N* = 0.472; (ii) *n*_2_*/N* = 0.472; *n*_3_*/N* = 0.056. (b) Leading edge as predicted by the three-species heterogeneous model, (*t*) (solid red), and the best-fit approximation given by the single-species homogeneous model, *S*(*t*) (blue dashed). (c) Error measure, 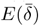, between the position of the leading edge given by the three-species heterogeneous model and the position predicted by the single-species homogeneous model as a function of cell size, 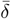. The black arrow denotes the best-fit value of cell size, 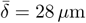. (d)-(e) Cell density profiles predicted by the three-species heterogeneous model, *𝒫*(*x, t*) (solid red), superimposed with density profiles given by the single-species homogeneous model calibrated with the best-fit value of *δ, P* (*x, t*) (solid blue). The continuum results for both models are presented at *t* = 0, 12, 24, 36, and 48 h. Black arrows denote the direction of increasing time. Results in (e) show a close-up comparison right near the leading edge, denoted by the gray shaded region in (d).

This manuscript is organised in the following way. In Section 2 we describe experimental data for a series of two-dimensional scratch assays that clearly involve a significant level of heterogeneity among the population. In Section 3 we introduce a mathematical model of the cell motility and adhesion. In particular, we focus on two analogues of the mathematical model: (i) a three-species heterogeneous model of cell motility where parameters including cell size, cell diffusivity and cell adhesion strength can vary between the subpopulations; and (ii) a more traditional single-species homogeneous model of cell motility where all cells in the population are treated as having the same cell size, cell diffusivity and cell adhesion strength. Results in Section 4 compare performance of the single-species homogeneous model as applied to data generated using the three-species heterogeneous model for different cell size distributions. Additional results presented in the Supplementary Material explore the role of: (i) heterogeneity in undirected (diffusive) migration, Set II; and (ii) heterogeneity in directed (adhesion/cell-to-cell contacts) migration, Set III. Finally, in Section 5 we summarise our result and propose potential extensions.

## 2 Experimental data

Monolayer scratch assays are performed using the IncuCyte ZOOM™ system (Essen BioScience). In all experiments we use the PC-3 prostate cancer cell line (Kaighn et al., 1979) from the American Type Culture Collection (ATCC™, Manassas, USA). After growing, cells are removed from the flask using TrypLE™ (ThermoFisher Scientific) in phosphate buffered saline, re-suspended in growth medium and seeded at a density of 20,000 cells per well in 96-well ImageLock plates (Essen BioScience). The diameter of each individual well is 9000 *µ*m.

Mitomycin-C is added at a concentration of 10 g/mL for two hours before a scratch is made in the monolayer of cells (Sadeghi et al., 1998). Mitomycin-C is a chemotherapy drug that blocks DNA replication and, consequently, stops proliferation. As a result of treatment the number of cells in the assay remains approximately constant since cells neither proliferate or die on the timescale of the experiment. Often scratch assays are performed using mitomycin-C treated cells so that the experiment focuses only upon the role of cell migration as opposed to the combined effects of cell migration and cell proliferation. A WoundMaker™ (Essen BioScience) is used to create identical scratches in the uniformly distributed populations. Medium is aspirated after scratching; each well is washed twice and refilled with fresh medium (100 *µ*L). Plates are incubated in the IncuCyte ZOOM™ and photographed every two hours for 48 hours. In total, these experiments are performed in eight of the 96 wells on the 96-well plate. In our work we use one of the experimental replicates at *t* = 0 h, shown in Figure 1, to quantify the heterogeneity in a cell population.

To quantify the heterogeneity in cell size we randomly select 184 cells from the experimental image in Figure 1(a) at *t* = 0 h. Assuming each cell can be treated as a disc, we estimate the equivalent diameter of each individual cell using the following approach. First, we use the histogram tool in Photoshop CS5 to count a number of pixels in the area occupied by each individual cell. The pixel count is converted to an area, *A*. Second, we estimate the equivalent diameter,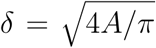 and use this data to produce histograms to illustrate and visualise the variability in cell size within the experiment. The resulting cell size distribution, presented as a histogram constructed with bin width 2.3 *µ*m, is shown in Figure 1(c). The bin width 2.3 *µ*m is chosen to demonstrate the fine structure within the cell population that is not normally incorporated in mathematical models of cell migration. However, the computational simulation of a population with the cell size distribution shown in Figure 1(c) is impractical since it would require significant computational resources to simulate the dynamics of 17 distinct subpopulations. As a compromise, we increase the bin width to reduce the number of distinct subpopulations while still retaining a sufficient number of bins to allow us to broadly characterise the heterogeneity in the population. Figure 1(d) demonstrates the histogram of cell sizes constructed using the same sample of cells with a larger bin size width of 15 *µ*m. Here, we have three subpopulations that capture the key trends in the heterogeneity in Figure 1(c) without needing to deal with 17 distinct subpopulations.

In this work we use experimental data to extract the cell size distribution at *t* = 0 h and use this data to generate the initial conditions in the three-species heterogeneous model (Set Ia, Figure 3). An interesting side effect of Mitomycin-C pretreatment is that cells increase in size abnormally fast compared to similar experiments without pretreatment (Matsiaka et al., 2018). As a result of pretreatment, the cell size distribution changes significantly with time, which, in turn, represents an additional degree of freedom in the problem. To keep our work tractable, we consider the most fundamental problem where we treat the cell size distribution as being constant through time, and we leave an extension to the case where the cell size distribution varies with time for future analysis.

## 3 Mathematical model

Discrete, stochastic models are often used to describe the spatial spreading of a population of cells, especially when the population of cells is not too large. Here, cells move and interact with each other via predefined force function, as illustrated schematically in Figure 2 (Newman and Grima, 2004; Callaghan et al., 2006; Hasenauer et al., 2011; Frascoli et al. 2013; Osborne et al., 2017). This approach is *individual-based* in the sense that knowledge about the movement of each individual is essential to infer the evolution of a density on the population-level scale. One of the most popular individual-based modelling approaches makes the assumption that the motion of each cell can be described by a Langevin stochastic differential equation (Newman and Grima, 2004; Middleton et al., 2014). As such, the system of *N* cells is described by a system of *N* stochastic differential equations of the form

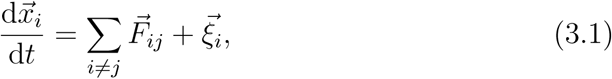

where 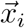 is the position vector of the *i*th cell, 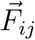 is the interaction force between cells *i* and *j*, and 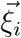 is the random stochastic force acting upon cell *i* (Middleton et al., 2014; George et al., 2017; Osborne et al., 2017). The interaction force, 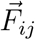, can be used to parametrise various features of cell populations, including heterogeneity. In fact, it is relatively straightforward to model heterogeneity in cell sizes in a discrete framework since the interaction force, 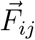, can be chosen to explicitly include the cell size as a parameter (Matsiaka et al., 2018). Here we can easily differentiate the population into an arbitrary number of subpopulations by assigning the value of the cell size to each member of the population. Despite the many advantages of this kind of individual-based modelling approach, such individual-based models are computationally inefficient as the number of cells, *N*, increases. This is because the computation time required to simulate such models increases with *N*.

In contrast, continuum models based on partial differential equations (PDEs) are much more convenient to model large cell populations because the time taken to solve continuum PDE models is independent of the size of the population (Sherratt and Murray, 1990; Sheardown and Cheng, 1995; Cai et al., 2007; Wise et al., 2008). Often, PDE models are derived using continuum-limit approximations of underlying discrete models and, as such, are able to retain certain features of a discrete model (Middleton et al., 2014; O’Dea and King, 2012). In this work we focus on a continuum model that is derived by taking the limit of a three-species heterogeneous individual-based model (Matsiaka et al., 2017). This approach allows us to conceptually incorporate key features of the heterogeneous cell populations into a discrete modelling framework, and then using a computationally efficient approach to solve the resulting continuum-limit PDE description of the underlying heterogeneous model.

We note that, due to the geometry of experiments presented in Figure 1, we are interested in the net movement of cells in only one direction, in this case the horizontal direction (Jin et al., 2016a). This is due to the fact that the net flux of cells in the vertical direction is, on average, zero because of the symmetry in the initial conditions of a scratch assay. Consequently, we focus on a one-dimensional continuum model and consider the evolution of the total cell population in the horizontal direction only. The use of a one-dimensional framework to describe two-dimensional scratch assays has been previously demonstrated to be a convenient approach to reduce the computational complexity while still describing the key features of the experiment (Matsiaka et al., 2018).

Here we employ a mean field model describing the spatial spreading of a population of cells composed of three distinct subpopulations. In one-dimension, the model can be written as

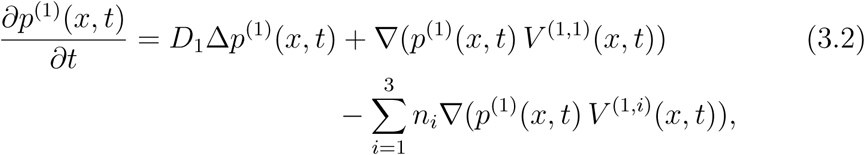

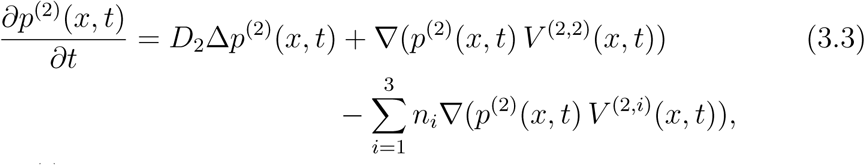

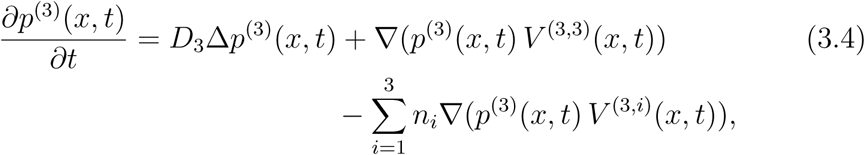

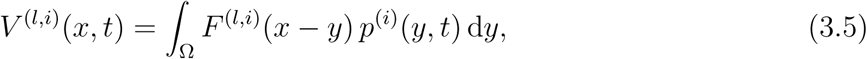

where *p*^(1)^(*x, t*), *p*^(2)^(*x, t*), and *p*^(3)^(*x, t*) are the cell densities associated with each subpopulation and depend on position *x* and time *t*. In this heterogeneous model, *D*_1_, *D*_2_, and *D*_3_ are diffusivities of subpopulations 1, 2, and 3, *n*_1_, *n*_2_, and *n*_3_ are the numbers of cells in each subpopulation, and *V* ^(*l,i*)^(*x, t*) is the velocity field of subpopulation *l* induced by subpopulation *i* (Matsiaka et al., 2017). The diffusivity constants parameterise the undirected migration of each subpopulation and the velocity fields describe the directed migration of each subpopulation that is driven by a combination of cell-to-cell adhesion and crowding effects.

The interaction force between subpopulations *l* and *i* that describes directed migration is given by

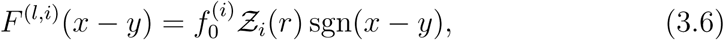

where 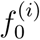is the dimensional amplitude of the interaction force acting on subpopulation 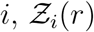 is a dimensionless function that parametrises different features of the cell-to-cell interactions, and *sgn* is the signum function. We choose to include long-range attraction that models cell-to-cell adhesion, and a short-range repulsion that reflects volume exclusion effects (Frascoli et al., 2013; Painter et al., 2010). A number of different phenomenological laws, 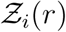, are used to model repulsive and adhesive intercellular forces (Murray et al., 2009; Jeon et al., 2010; Middleton et al., 2014). In our work we adopt modified Morse potential in the form

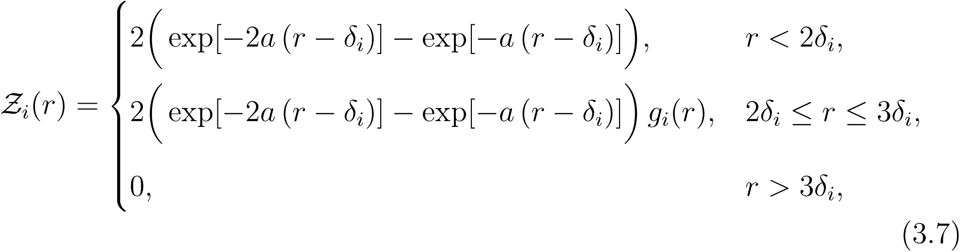

where *a* is the parameter that controls the shape of the force function, *δ*_*i*_ is the cell size in the subpopulation *i, i* = 1, 2, 3, and *r* = |*x* − *y*|. We fix the value of the shape parameter at *a* = 0.08 *µ*m^−1^ (Matsiaka et al., 2017). The function 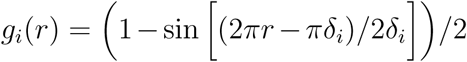 is the Tersoff cut-off function introduced to impose a finite range of intercellular interactions (Tersoff, 1988). A sketch of the potential function given by Equation (3.7) for different values of the parameter *a* is shown in Figure 2(b) confirming that this potential function describes short range repulsion, longer range attraction and no interactions at over much longer distances. In summary, the key parameters in the heterogeneous three-species model are: (i) the cell sizes, *δ*_1_, *δ*_2_ and *δ*_3_; (ii) the cell diffusivities, *D*_1_, *D*_2_ and *D*_3_; and (iii) the amplitudes of interaction forces, 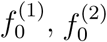 and 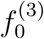. In this work we will systematically explore how heterogeneity in each of these three key parameters influences whether we need to consider a complex heterogeneous multi-species model or whether we can describe the spatial spreading of a cell population using relatively simple homogeneous, single-species models. Since our experimental data in Figure 1 allows us to explicitly characterise the heterogeneity in cell size, all results in the main document focus on cell size. Additional results in the Supplementary Material focus on heterogeneity in diffusivity and amplitude of interaction forces to provide additional insight into the role of heterogeneity in these kinds of experiments.

We define the total density of the heterogeneous population as

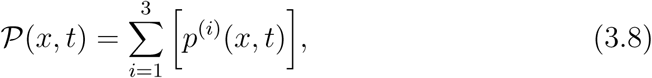

where *p*^(*i*)^(*x, t*) is the cell density of subpopulation *i* = 1, 2, 3 predicted by Equations (3.2)-(3.4), and *𝒫* (*x, t*) is the total cell density. It is important to interpret the solutions of Equations (3.2)-(3.4) in terms of total cell density since standard experimental protocols do not normally facilitate the measurement of spatial and temporal distributions of various subpopulations (Cai et al., 2007; Treloar et al., 2014).

We can reduce the three-species heterogeneous system of equations, Equations (3.2)-(3.4), to obtain a single-species homogeneous model in the form,

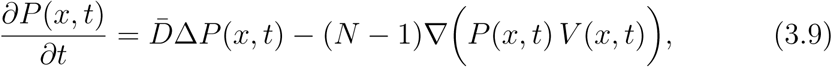

where *P* (*x, t*) is the cell density of the total population, 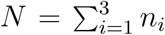 is the total number of cells in the population. Here we assume that the cell size, diffusivity and strength of the interaction force for each population is constant, giving 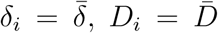, and 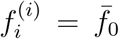 for *i* = 1, 2, 3. The key differences between the homogeneous single-species model, Equation (3.9), and the three-species heterogeneous model, Equations (3.2)-(3.4) are: (i) the three-species heterogeneous model incorporates three advection-diffusion equations while the single-species homogeneous model is given by a single advection-diffusion equation; (ii) the three-species heterogeneous model contains up to nine free parameters as opposed to three parameters in the single-species homogeneous model.

The initial conditions in all simulations are chosen to mimic a cell front, such as that shown in our experimental data set, Figure 1(a). As such, we adopt an initial cell distribution in the form of the one-dimensional step function,

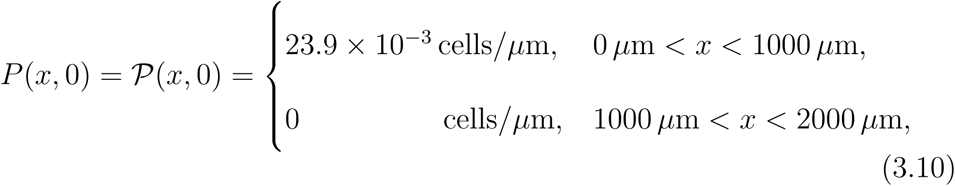

on 0 < *x* < 2000 *µ*m, which is consistent with a length-scale of a typical *in vitro* experiment (Jin et al., 2016a). The initial cell distribution in the heterogeneous model is given by the sum of initial densities of three subpopulations, 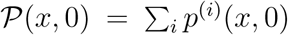, where the density of each subpopulation, *p*^(*i*)^(*x,* 0), varies between each cell size distribution and can be inferred from the histograms in Figure 3(a). The value of the initial density of the total population is chosen to represent fairly confluent population of cells. For example, the simulation of the three-species population with the monotonically decreasing cell size distribution, Set Ia, is initiated with the confluence level of approximately 65% of maximum packing density, which is fairly typical for scratch assay experiments (Jin et al., 2016; Matsiaka et al., 2017). We note that the boundary of the experimental image in Figure 1(a) is not a physical boundary and cells can freely move across this boundary because the image captures only a small fraction of a much larger experimental domain (Simpson et al., 2018). During the experiment, cells freely migrate, in each direction, across the boundary. However, since the density of cells away from the scratch is spatially uniform, the net flux of cells across the boundary of the image is zero. To capture this situation we impose zero net flux boundary conditions at *x* = 0 *µ*m and *x* = 2000 *µ*m.

All continuum results for single-species homogeneous and three-species heterogeneous models, given by Equation (3.9) and Equations (3.2)-(3.4), respectively, are solved numerically using the method of lines with Δ*x* = 4 *µ*m and Δ*t* = 0.005 h on 0 < *x* < 2000 *µ*m (Matsiaka et al., 2017). We find that this choice of spatial and temporal discretisations are sufficiently fine to produce grid independent results. The detailed discretisation scheme used in this work is presented in the Supplementary Material.

## 4 Results and Discussion

To investigate the ability of a single-species homogeneous model to capture the behaviour of the three-species heterogeneous analogue, we consider a series of case studies. In these case studies we vary only one parameter at a time to simplify our analysis and to focus on the impact of each individual parameter. Another approach would be to use the mathematical models to explore heterogeneity multiple parameter at the same time. However, in this first instance, we prefer to take a more fundamental approach and examine the role of heterogeneity in each parameter separately. In the first set of experiments, Set I, we vary the cell size, 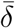, while keeping 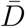 and 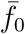 fixed at 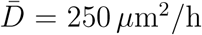 and 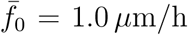. The values of *D*_*i*_ and 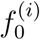 in the heterogeneous three-species model are fixed at *D*_*i*_ = 250 *µ*m^2^*/*h and 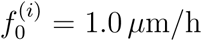 for *i* = 1, 2, 3. These values of diffusivity and amplitude of cell-to-cell interaction forces are based on detailed experimental measurements reported previously (Matsiaka et al., 2019).

There are number of ways to quantify performance of the single-species homogeneous model in our framework. The position of the leading edge of the spreading population is routinely used by experimentalists to provide quantitative insights into the rate of spatial spreading of a cell population (Treloar and Simpson, 2013; Johnston et al., 2014; Kollimada et al., 2016; Nardini et al., 2016; Bobadilla et al., 2019). Therefore, we quantify the discrepancy between the solution of the heterogeneous three-species model and the homogeneous single-species model using an error measure, 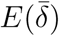, associated with the position of the leading edge,

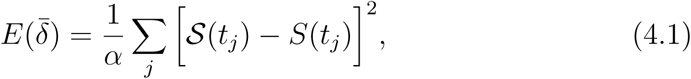

where *S*(*t*_*j*_) is the position of the leading edge according to the three-species heterogeneous model at time *t*_*j*_, *S*(*t*_*j*_) is the position of the leading edge predicted by the single-species homogeneous model, and *α* = 49 is the number of discrete time points we use to compute 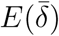. In both scenarios the position of the leading edge is computed as the coordinate on the one-dimensional domain where the density is 1% of the initial density (Treloar and Simpson, 2013). An alternative approach is to use an error measure based on the discrepancy between cell density profiles. At first, this approach of using the entire cell density profile might be thought to be preferable to working with leading edge data since density profiles incorporate much more detailed spatial information than just using the position of the leading edge. However, extracting the density data from experiments is much more tedious because it often involves manual cell counting in regions where cell densities are high and this is both difficult to reproduce and very time consuming (Treloar et al. 2014). Therefore, to keep our work as practical as possible, here we present only results with an error measure solely based on the leading edge data. Additional result that measure the discrepancy between the models using the entire density information are presented in the Supplementary Material (Figure A.1 and Figure A.2), and we find that this more complicated approach gives very similar results to the leading edge data. Therefore, in this work, we focus on the using leasing edge data.

The experimental distribution of cell sizes in Figure 1(d) provides insights into potential choices of the cell size distribution in Equations (3.2)-(3.4). Here we define three subpopulations based on the equivalent cell size: small (*δ*_1_ = 18 *µ*m), medium (*δ*_2_ = 34 *µ*m), and large cells (*δ*_3_ = 50 *µ*m). For simplicity, we set the fractions of small and medium cells to be equal and refer to this distribution as a monotonically decreasing distribution of cell sizes (Set Ia, Figure 3). After considering the experimentally-motivated monotonically decreasing distribution, we then systematically explore: (i) uniform (Set Ib, Figure 4), non-monotonic (Set Ic, Figure 5), and (iii) monotonically increasing distributions (Set Id, Figure 6).

**Fig. 4.**
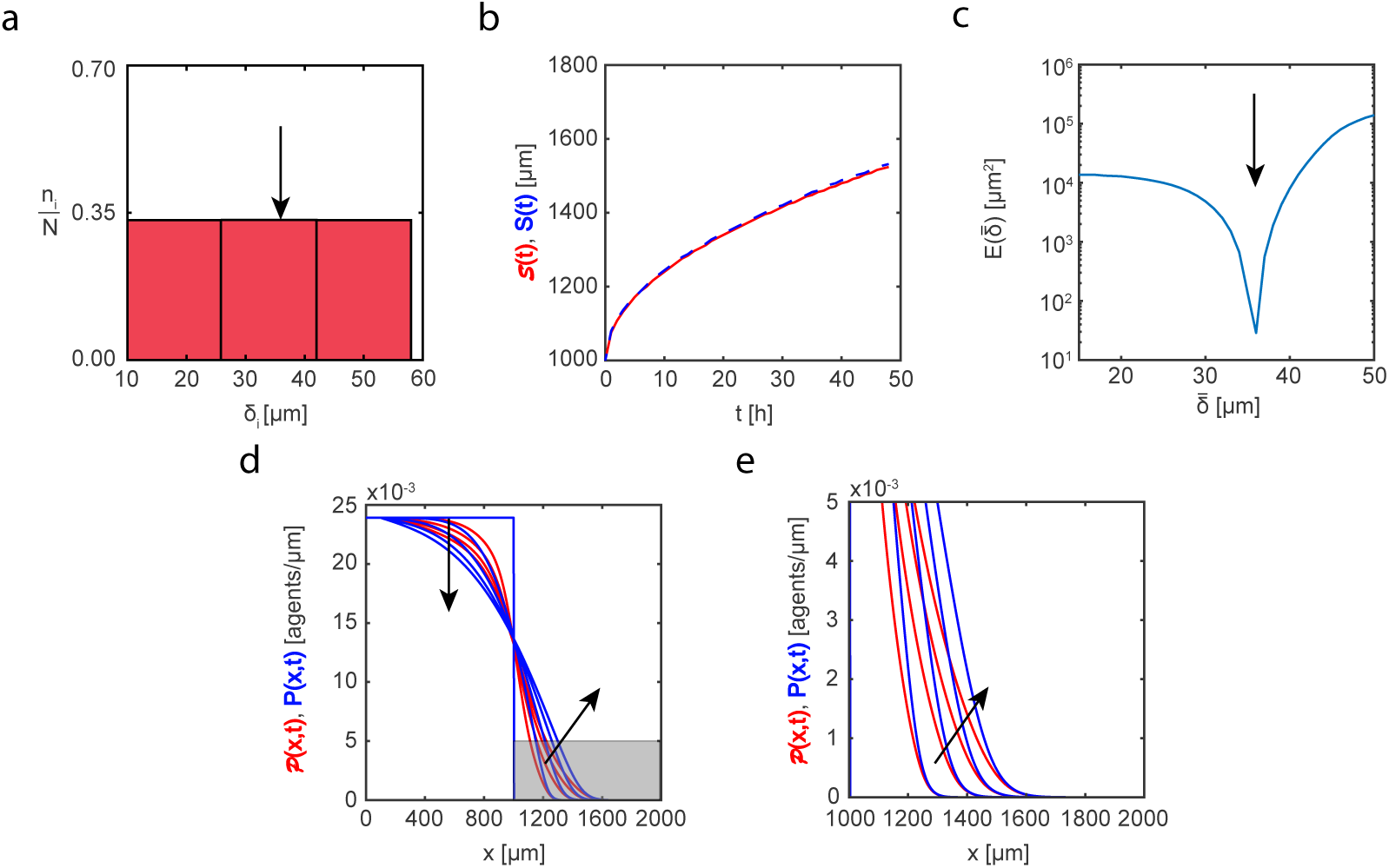
Set Ib. Heterogeneity in cell sizes: uniform distribution. (a) Cell size distribution adopted in the three-species heterogeneous model, Equations (3.2)-(3.4). Here the proportions of cells of different sizes are set to: (i) *n*_1_*/N* = 0.33(3); (ii) *n*_2_*/N* = 0.33(3); (iii) *n*_3_*/N* = 0.33(3). (b) Leading edge as predicted by the three-species heterogeneous model, (*t*) (solid red), and the best-fit approximation given by the single-species homogeneous model, *S*(*t*) (blue dashed). (c) Error measure, 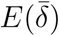, between the position of the leading edge given by the three-species heterogeneous model and the position predicted by the single-species homogeneous model as a function of cell size, 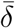. The black arrow denotes the best-fit value of cell size, 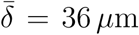. (d)-(e) Cell density profiles predicted by the three-species heterogeneous model, 𝒫 (*x, t*) (solid red), superimposed with density profiles given by the single-species homogeneous model calibrated with the best-fit value of 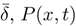 (solid blue). The continuum results for both models are presented at *t* = 0, 12, 24, 36, and 48 h. Black arrows denote the direction of increasing time. Results in (e) show a close-up comparison right near the leading edge, denoted by the gray shaded region in (d).

**Fig. 5.**
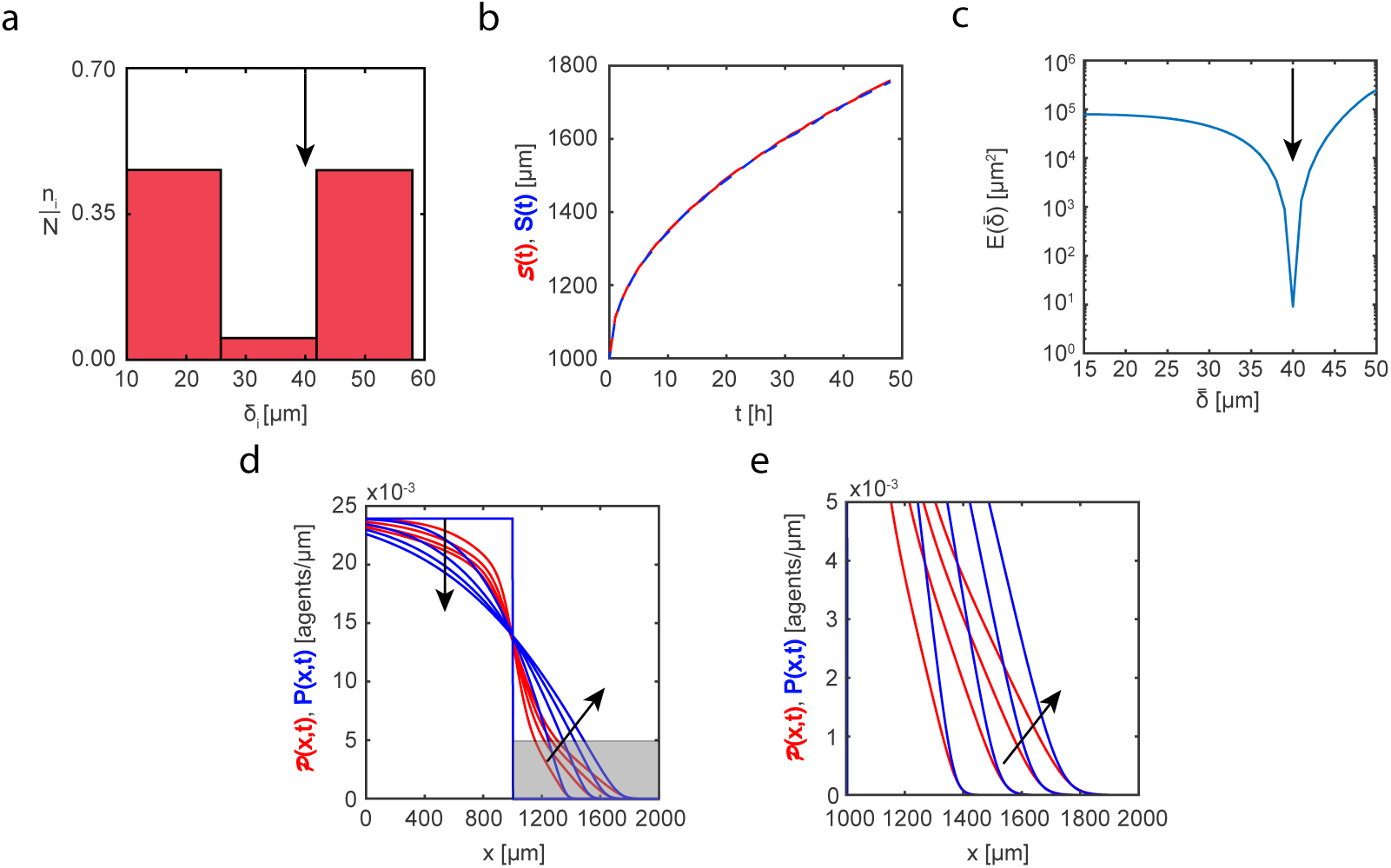
Set Ic. Heterogeneity in cell sizes: non-monotonic distribution. (a) Cell size distribution adopted in the three-species heterogeneous model, Equations (3.2)-(3.4). Here the proportions of cells of different sizes are set to: (i) *n*_1_*/N* = 0.472; (ii) *n*_2_*/N* = 0.056; (iii) *n*_3_*/N* = 0.472. (b) Leading edge as predicted by the three-species heterogeneous model, (*t*) (solid red), and the best-fit approximation given by the single-species homogeneous model, *S*(*t*) (blue dashed). (c) Error measure, 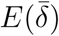, between the position of the leading edge given by the three-species heterogeneous model and the position predicted by the single-species homogeneous model as a function of cell size, 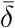. The black arrow denotes the best-fit value of cell size, 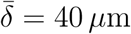. (d)-(e) Cell density profiles predicted by the three-species heterogeneous model, 𝒫(*x,t*) (solid red), superimposed with density profiles given by the single-species homogeneous model calibrated with the best-fit value of 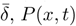 (solid blue). The continuum results for both models are presented at *t* = 0; 12; 24; 36; and 48 h. Black arrows denote the direction of increasing time. Results in (e) show a close-up comparison right near the leading edge, denoted by the gray shaded region in (d).

**Fig. 6.**
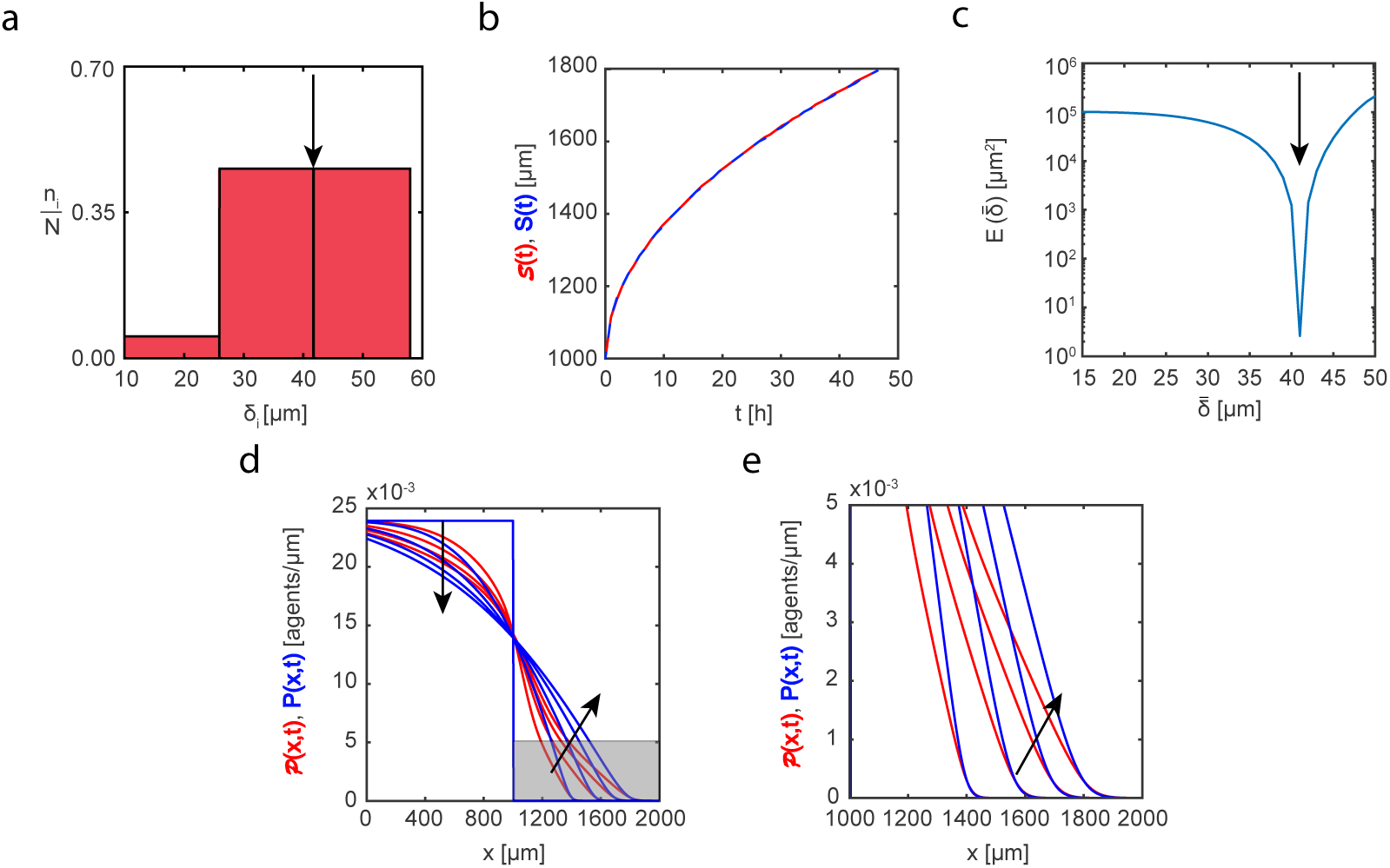
Set Id. Heterogeneity in cell sizes: monotonically increasing distribution. (a) Cell size distribution adopted in the three-species heterogeneous model, Equations (3.2)-(3.4). Here the proportions of cells of different sizes are set to: (i) *n*_1_*/N* = 0.056; (ii) *n*_2_*/N* = 0.472; (iii) *n*_3_*/N* = 0.472. (b) Leading edge as predicted by the three-species heterogeneous model, 𝓢(*t*) (solid red), and the best-fit approximation given by the single-species homogeneous model, *S*(*t*) (blue dashed). (c) Error measure, 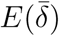, between the position of the leading edge given by the three-species heterogeneous model and the position predicted by the single-species homogeneous model as a function of cell size, 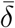. The black arrow denotes the best-fit value of cell size, 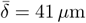. (d)-(e) Cell density profiles predicted by the three-species heterogeneous model, 𝒫(*x,t*) (solid red), superimposed with density profiles given by the single-species homogeneous model calibrated with the best-fit value of 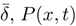 (solid blue). The continuum results for both models are presented at *t* = 0; 12; 24; 36; and 48 h. Black arrows denote the direction of increasing time. Results in (e) show a close-up comparison right near the leading edge, denoted by the gray shaded region in (d).

Figure 3(b) compares the leading edge prediction, *S*(*t*), given by the three-species heterogeneous model with the associated best-fit match, *S*(*t*), predicted by the single-species homogeneous model. Our systematic computation of the error measure, 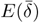, demonstrates a clear minimum which ensures the unique choice of a best-fit cell size, 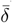. Results in Figure 3(d) superimposes the solution of the three-species heterogeneous model with the solution of the single-species homogeneous model parameterised with the best fit cell size. Comparing the time evolution of the spreading density profiles in Figure 3(d) (with additional details at the leading edge shown in the magnified region in Figure 3(e)) we see that the appropriately parameterised single-species homogeneous model captures the temporal evolution of the spreading profile given by the heterogeneous model remarkably accurately. In particular, the density profiles predicted by the single-species homogeneous model match both the position and shape of the density profiles generated by the three-species heterogeneous model. These results imply that in this case it would be reasonable to use a much simpler single-species homogeneous model to describe and predict this spatial spreading.

Visual inspection of the results in Figures 3 - 6 suggests that we can always find a unique, well-defined value of the cell size in the single-species homogeneous model to provide an accurate prediction of the temporal evolution of the position of a leading edge of the spreading heterogeneous cell populations regardless of the underlying cell size distribution in the three-species heterogeneous model (Figures 3(b)-6(b)). In contrast, the quality of match between the shape of the density profiles for the three-species heterogeneous model and the single-species homogeneous model varies significantly between different cell size distributions. For example, the experimentally motivated distribution in Figure 3(a) (Set Ia) leads to a remarkably good match between the three-species heterogeneous model and the single-species homogeneous model. Similarly, the uniform distribution shown in Figure 4(a) (Set Ib) also leads to a reasonably good quality of match between two different models. In contrast, the density profiles associated with the non-monotonic cell size distribution (Figure 5, Set Ic) and monotonically increasing cell size distribution (Figure 6, Set Id) show a relatively poor match. In these cases, it would seem prudent not to use a simpler single-species homogeneous model to simulate and predict these experiments.

The values of the cell size, 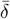, that produce best match between the single-species homogeneous and three-species heterogeneous models vary significantly between different cell size distributions. For example, the best-fit value of the cell size for the uniform distribution (Figure 4, Set Ib), 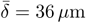, is quite close to the weighted average value of 34 *µ*m for the distribution in Figure 4(a). This indicates that the choice of a simple weighted average of the cell sizes might be a reasonable way to to parameterise the single-species homogeneous model if the experimentally observed distribution is close to uniform. We observe similar agreement for best-fit values of the cell size in the case of monotonically decreasing (Set Ia) and monotonically increasing (Set Id) cell size distributions, shown in Figure 3 and Figure 6, respectively. In contrast, the best-fit value of the cell size for the non-monotonic distribution (Set Ic), 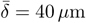, differs significantly from the weighted average of 34 *µ*m. Therefore, these results suggest that great care ought to be exercised when taking a distribution of parameter values and attempting to select the most appropriate single representative value of that parameter.

In addition to the results in Figures 3 - 6 exploring the role of heterogeneity in cell size, we present an additional suite of results where we systematically explore the role of heterogeneity in diffusivity (Set II) and amplitude of interaction forces (Set III) while keeping the cell size constant in all subpopulations. These additional results are presented in the Supplementary Material document. Both Set II and Set III data sets demonstrate exceptional quality of match between the three-species heterogeneous simulation data and its best-fit single-species homogeneous equivalent. Again, these additional results provide guidance about when it is reasonable to approximate a more complicated heterogeneous mathematical model with a simpler single-species homogeneous model.

## 5 Conclusions

In this work, we explore the role of heterogeneity in the context of studying how an initially confined population of cells can spread into surrounding initially unoccupied regions, as in the case of a scratch assay. We use a three-species heterogeneous model of cell motility, account for undirected cell motility, short range repulsion (crowding) and longer range adhesion, to capture experimentally observed heterogeneity in cell sizes from a new experimental data set from a two-dimensional scratch assay as shown in Figure 1. Our continuum models account for the undirected random motility, cell-to-cell adhesion, and cell crowding. The single-species homogeneous model is applied to each set of three-species heterogeneous simulation data in an attempt to match cell density profiles.

To analyse the performance of the single-species homogeneous model to capture data from our three-species heterogeneous model we consider four different cell size distributions: (i) monotonically decreasing distribution, (ii) uniform distribution, (iii) non-monotonic distribution, and (iv) monotonically increasing distribution. Overall, for a set of experimentally-motivated parameter combinations, we find that the standard single-species homogeneous model is able to accurately predict the position of the leading edge for all case studies presented. However, the quality of the match between the shape of the density profiles varies significantly depending on the details of the form of the heterogeneity present. For example, the monotonically decreasing distribution (Set Ia) demonstrates remarkable goodness of fit between the two sets of density profiles, as shown in Figure 3(d). This result is important because the monotonically decreasing cell size distribution is chosen to mimic the distribution of the cell sizes observed in our new experimental data set, shown in Figure 1. Similarly, the homogeneous distribution, Figure 4, shows that single-species homogeneous model is able to accurately replicate the three-species heterogeneous model results. This is an expected result because in this special case the cells of each subpopulation are the same size. In contrast, the single-species homogeneous model does not perform so well when applied to both non-monotonic and monotonically increasing distributions in Figures 5-6, respectively. Additionally we explore potential heterogeneity in diffusivity and amplitude of the cell-to-cell interactions (Supplementary Material). Overall, our results suggest that for certain cell size distributions, a simple and computationally efficient single-species homogeneous model is preferable over a thee-species heterogeneous model.

There are number of ways this work can be extended which we leave for future analysis. All our simulations and analysis focus on treating the heterogeneity in the population of cells by considering the total population to be composed of three distinct subpopulations. For more extreme forms for heterogeneity, such as multi-modal distributions, the results presented in this work could be extended by considering additional subpopulations. Another simplification that we invoke is to assume that the measured heterogeneity remains constant for the duration of the experiment. Future studies could address the significantly more complicated question of allowing the distributions to evolve in time on the same time scale as the experiment to see if it is still possible to use a simpler homogeneous model in this more complicated scenario. Another avenue for further exploration would be to consider heterogeneity in more than one parameter at a time, whereas in this work we have taken the most fundamental approach and examined heterogeneity in just one parameter in isolation from the others. For both of these extensions, the modelling framework presented in this study can be extended to explore these additional features, and we leave such extensions for future consideration. Another option for extending the work would be to consider further details in the mathematical models, such as the effects of combined cell migration and combined cell proliferation. Here we have not pursued this approach because our experimental data set has been carefully prepared to exclude the effects of proliferation so that we can focus just on cell migration and heterogeneity in cell migration alone.

## Acknowledgements

This work is supported by the Australian Research Council (DP170100474). Computational resources are provided by the High Performance Computing and Research Support Group at QUT. REB is a Royal Society Wolfson Research Merit Award holder, would like to thank the Leverhulme Trust for a Research Fellowship and also acknowledges the BBSRC for funding via grant no. BB/R000816/1. We appreciate helpful comments provided by two anonymous referees.

