## Supplementary Material for "Continuum descriptions of spatial spreading for heterogeneous cell populations: theory and experiment"

---

**Contents**

A Heterogeneity in the cell size assessed using density profile data 2

B Set II: Heterogeneity in the interaction forces 6

C Set III: Heterogeneity in the diffusivity 11

D Discretisation scheme for the single-species homogeneous model and heterogeneous three-species model 16

* Corresponding author

### A Heterogeneity in the cell size assessed using density profile data

In this section we repeat the analysis contained in Figure 3 and 5 (Main document), except that we use a different measure of the discrepancy, that is instead of using the leading edge data, we use a measure that is based on the entire cell density profile. In this analysis we keep $D_i$ and $f_0^{(i)}$ fixed at $D_i = 250\,\mu\text{m}^2/\text{h}$ and $f_0^{(i)} = 1.0\,\mu\text{m/h}$ for $i = 1, 2, 3$, respectively (Set I, Main document). The main difference is that here we define the error measure, $\mathbb{E}(\bar{\delta})$, as the mean square difference between the density profiles given by the three-species heterogeneous model and profiles predicted the single-species homogeneous model,

$$\mathbb{E}(\bar{\delta}) = \frac{1}{\alpha \mathcal{N}} \sum_i \sum_j \Big[\mathcal{P}(x_i, t_j) - P(x_i, t_j)\Big]^2, \tag{A.1}$$

where $\mathcal{P}(x_i, t_j)$ is the total agent density given by the three-species heterogeneous model, $P(x_i, t_j)$ is the agent density predicted by the single-species homogeneous model, $\alpha = 49$ is the number of discrete time points that we used to compute $\mathbb{E}(\delta)$, and $\mathcal{N} = 500$ is the number of discrete spatial points in the discretisation scheme.

Results in Figure A1 and A2 are analogous to those results in Figure 3 and 5 (Main document). The best-fit value of the cell size, $\bar{\delta} = 26\,\mu$m, is relatively close to a best-fit estimate of $\bar{\delta} = 28\,\mu$m reported in the main document using the error measure, $E(\bar{\delta})$, based on the position of the leading edge (compare Figure A.1(b) and Figure 3(c) in the main document). Here we find that the single species homogeneous model can be used to accurately describe the three-species heterogeneous data for both error measures, and this is obvious when we visually compare Figure A.1(c) and Figure 3(d) in the main document. In contrast, the density profiles associated with the best-fit homogeneous model in Figure A.2(c) do not provide an accurate match the three-species heterogeneous data generated. This means that the heterogeneous density profiles associated with the non-monotonic distribution cannot be faithfully replicated using a simpler homogeneous model. In summary, the conclusions we draw in the main document about the suitability of the homogeneous model to accurately describe results generated using the heterogeneous model are the same regardless of whether we use leading edge data or the entire cell density profile to measure the discrepancy between the two modelling frameworks.

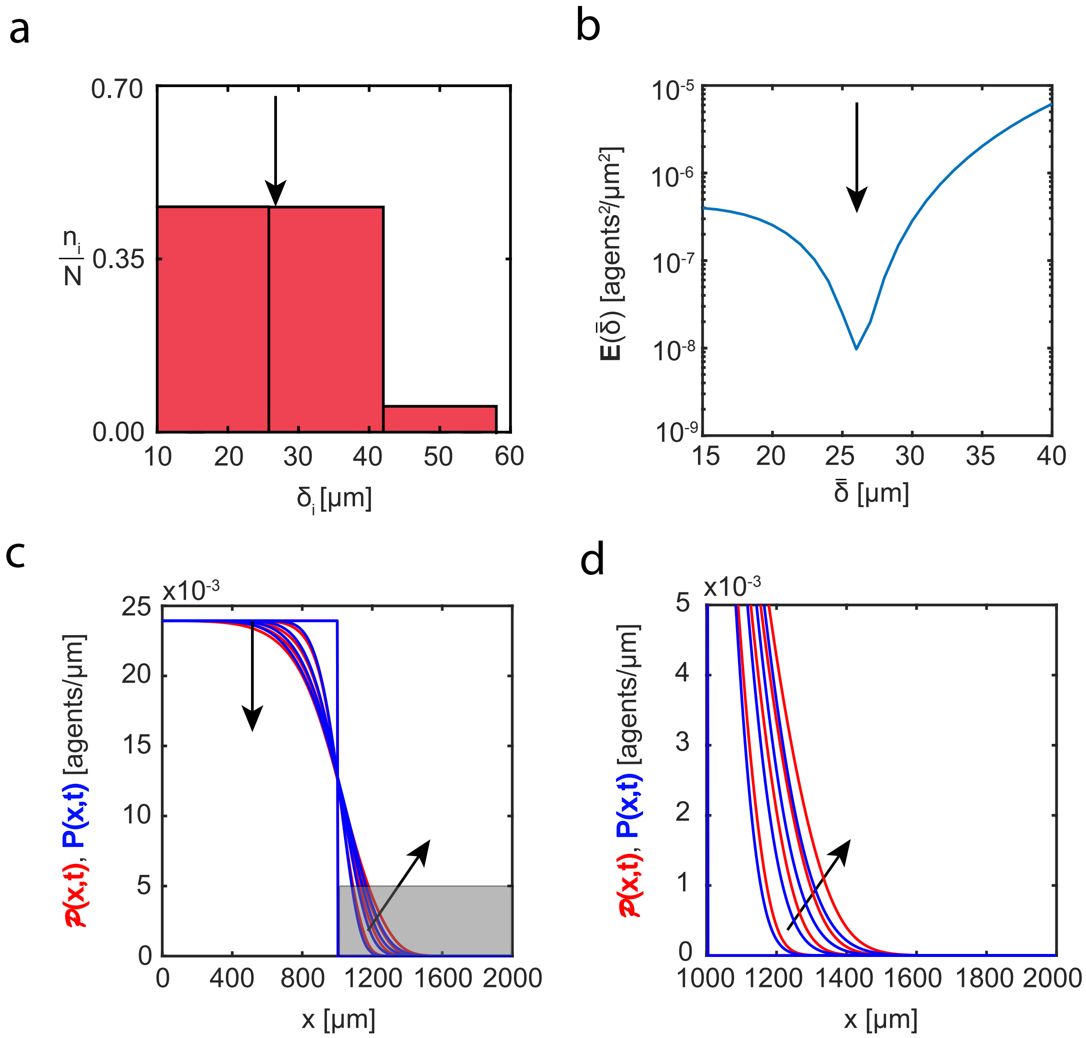


Fig. A.1. Heterogeneity in cell sizes: monotonically decreasing distribution. (a) Cell size distribution adopted in the three-species heterogeneous model, Equations (3.2)-(3.4), (main document). Here the proportions of cells of different sizes are set to: (i) $n_1/N = 0.472$; (ii) $n_2/N = 0.472$; (iii) $n_3/N = 0.056$. (b) Error measure, $\mathbb{E}(\bar{\delta})$, between the cell density profiles, $\mathcal{P}(x,t)$, given by the three-species heterogeneous model and density profiles, $P(x,t)$, predicted by the single-species homogeneous model as a function of cell size, $\bar{\delta}$. The black arrow denotes the best-fit value of cell size, $\bar{\delta} = 26\,\mu$m. (c)-(d) Cell density profiles predicted by the three-species heterogeneous model, $\mathcal{P}(x,t)$ (solid red), superimposed with density profiles given by the single-species homogeneous model calibrated with the best-fit value of $\bar{\delta}$, $P(x,t)$ (solid blue). The continuum results for both models are presented at $t = 0, 12, 24, 36,$ and $48\,$h. Black arrows denote the direction of increasing time. Results in (d) show a close-up comparison right near the leading edge, denoted by the gray shaded region in (c).

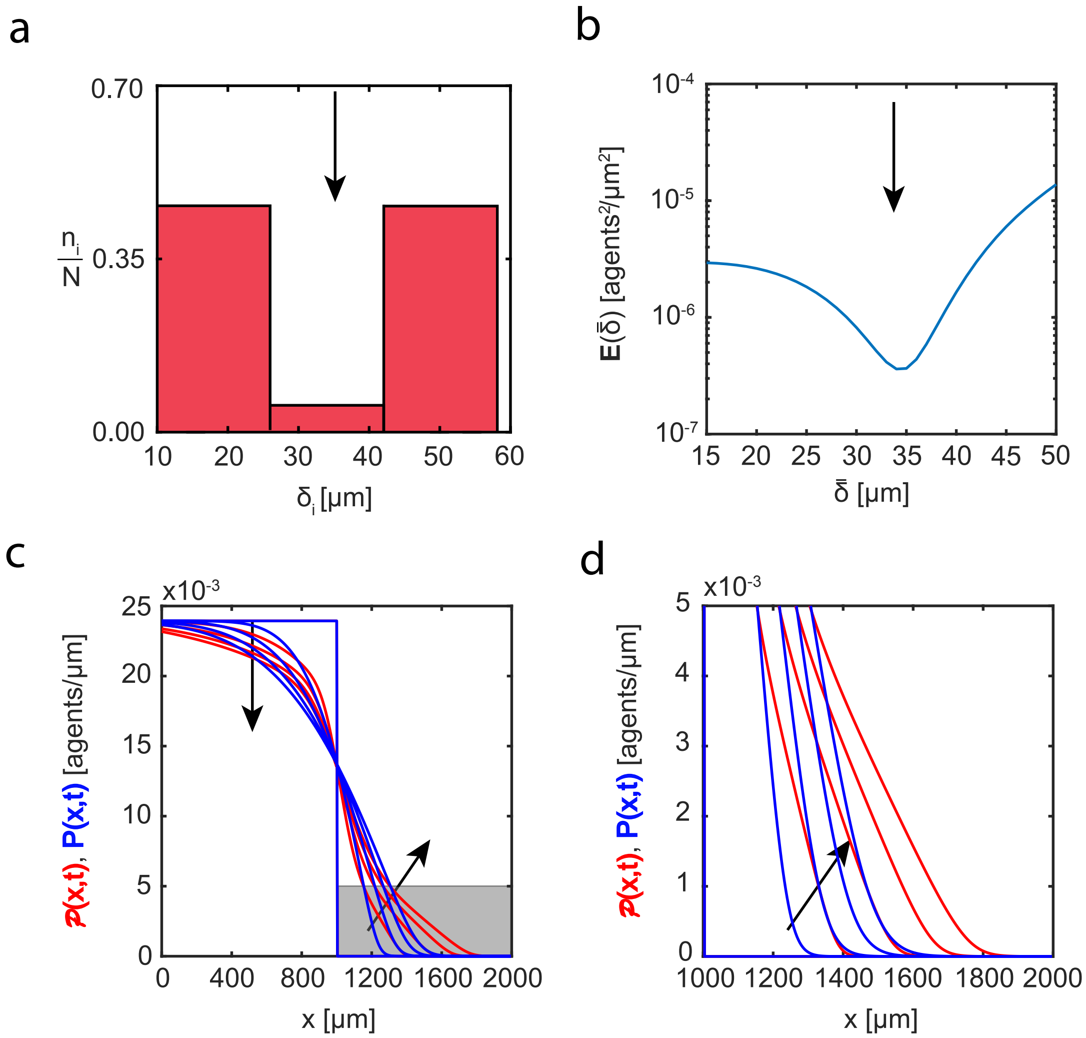


Fig. A.2. Heterogeneity in cell sizes: non-monotonic distribution. (a) Cell size distribution adopted in the three-species heterogeneous model, Equations (3.2)-(3.4), (main document). Here the proportions of cells of different sizes are set to: (i) $n_1/N = 0.472$; (ii) $n_2/N = 0.056$; (iii) $n_3/N = 0.472$. (b) Error measure, $\mathbb{E}(\bar{\delta})$, between the cell density profiles, $\mathcal{P}(x,t)$, given by the three-species heterogeneous model and density profiles, $P(x,t)$, predicted by the single-species homogeneous model as a function of cell size, $\bar{\delta}$. The black arrow denotes the best-fit value of cell size, $\bar{\delta} = 34\,\mu$m. (c)-(d) Cell density profiles predicted by the three-species heterogeneous model, $\mathcal{P}(x,t)$ (solid red), superimposed with density profiles given by the single-species homogeneous model calibrated with the best-fit value of $\bar{\delta}$, $P(x,t)$ (solid blue). The continuum results for both models are presented at $t = 0, 12, 24, 36$, and 48 h. Black arrows denote the direction of increasing time. Results in (d) show a close-up comparison right near the leading edge, denoted by the gray shaded region in (c).

### B Set II: Heterogeneity in the interaction forces

In this data set we explore the heterogeneity in the interaction forces where we fix values of the diffusivity, $D_i = 250\,\mu\text{m}^2/\text{h}$, and the cell size, $\delta_i = 34\,\mu\text{m}$ for $i = 1, 2, 3$. We note that these estimates are typical parameter values for PC-3 cells (Matsiaka et al., 2019). To analyse performance of the single-species homogeneous model (Equation (3.9), main document) applied to data generated by the three-species heterogeneous model (Equations (3.2)-(3.4), main document) we consider four interaction force distributions: (i) uniform distribution, Figure B.1(a), (ii) monotonically decreasing distribution, Figure B.2(a), (iii) non-monotonic distribution, Figure B.3(a), and (iv) monotonically increasing distribution, Figure B.4(a). For all cases presented we are able to predict a position of the leading edge as well as accurately describe cell density profiles given by the three-species heterogeneous model.

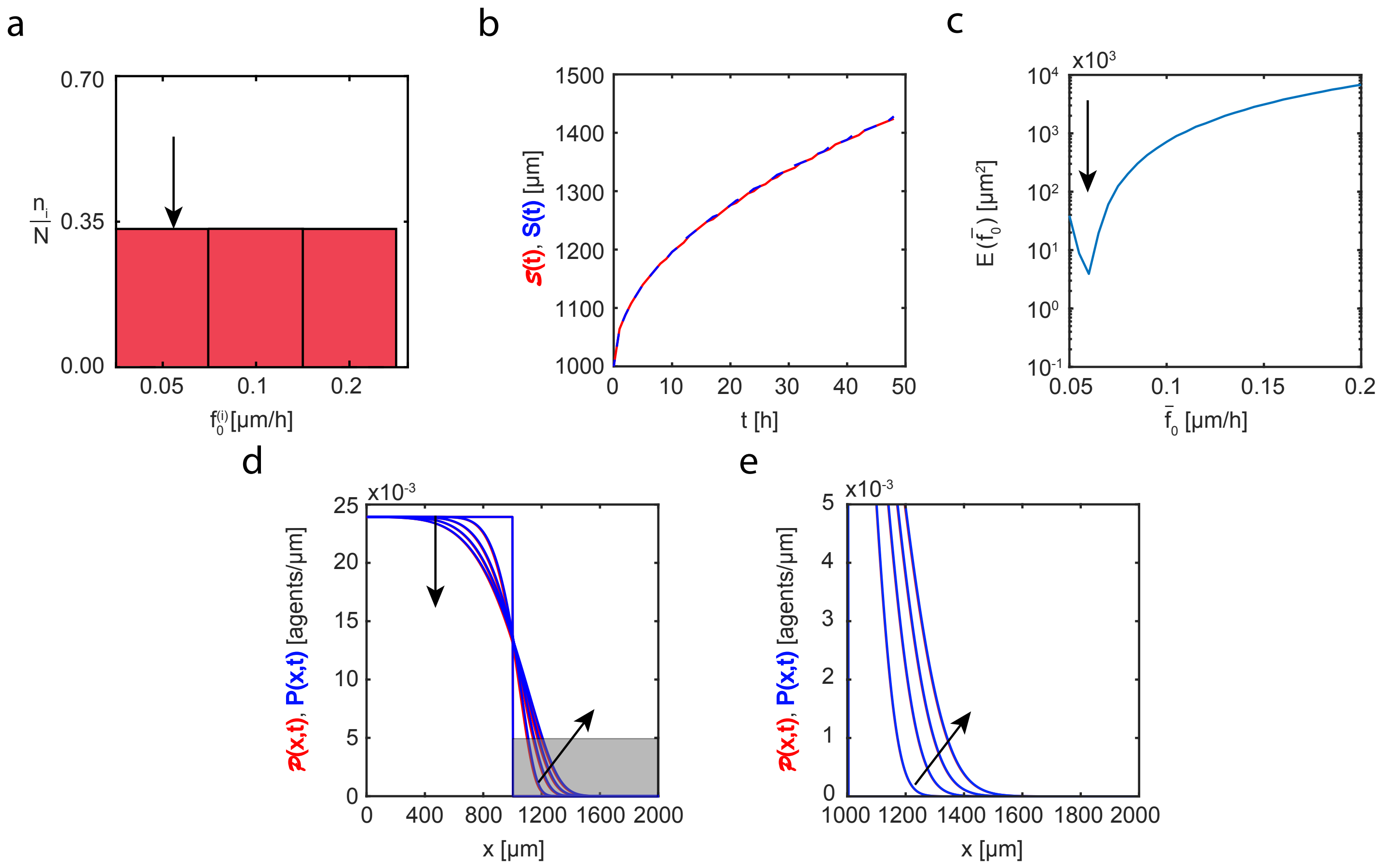


Fig. B.1. Set IIa. Heterogeneity in interaction forces: uniform distribution. (a): Interaction force distribution adopted in the three-species heterogeneous model, Equations (3.2)-(3.4) (main document). Here the proportions of cells of different sizes are set to: (i) $n_1/N = 0.333$; (ii) $n_2/N = 0.333$; (iii) $n_3/N = 0.333$. (b): Leading edge predicted by the three-species heterogeneous model, $\mathcal{S}(t)$ (solid red), and the best-fit approximation given by the single-species homogeneous model, $S(t)$ (blue dashed). (c): Error measure, $E(\bar{f}_0)$, between the position of the leading edge given by the three-species heterogeneous model and the position predicted by the single-species homogeneous model as a function of amplitude of the interaction force, $\bar{f}_0$. (d)-(e): Cell density profiles predicted by the three-species heterogeneous model, $\mathcal{P}(x,t)$ (solid red), superimposed with density profiles given by the single-species homogeneous model calibrated with the best-fit value of $\bar{f}_0$, $P(x,t)$ (solid blue). The black arrow denotes the best-fit value of interaction force, $\bar{f}_0 = 0.055\,\mu\text{m/h}$. The continuum results for both models are presented at $t = 0, 12, 24, 36,$ and $48\,\text{h}$. Black arrows denote the direction of increasing time. Results in (e) show a close-up comparison right near the leading edge, denoted by the gray shaded region in (d).

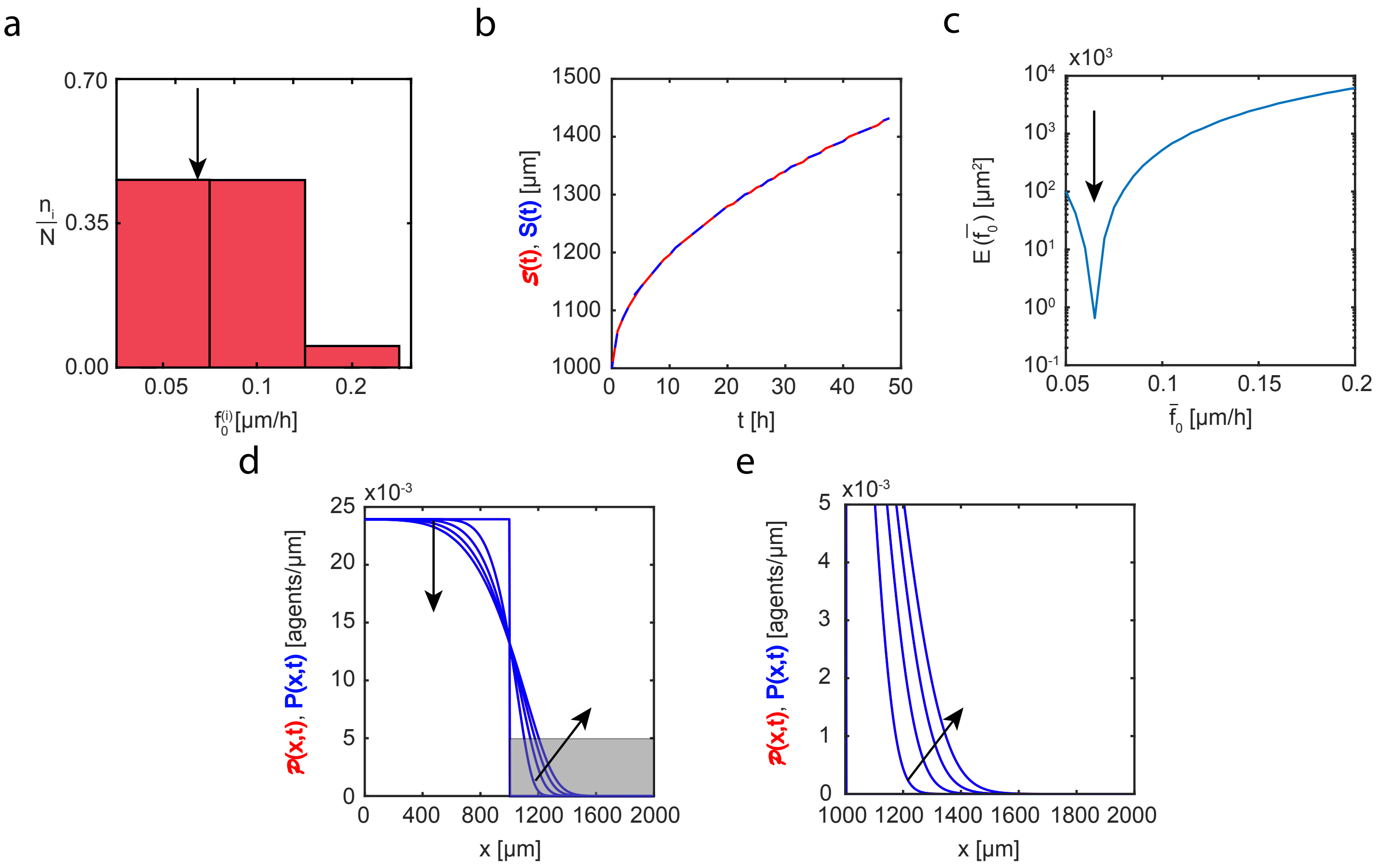


Fig. B.2. Set IIb. Heterogeneity in interaction forces: monotonically decreasing distribution. (a): Interaction force distribution adopted in the three-species heterogeneous model, Equations (3.2)-(3.4) (main document). Here the proportions of cells of different sizes are set to: (i) $n_1/N = 0.472$; (ii) $n_2/N = 0.472$; (iii) $n_3/N = 0.056$. (b): Leading edge predicted by the three-species heterogeneous model, $\mathcal{S}(t)$ (solid red), and the best-fit approximation given by the single-species homogeneous model, $S(t)$ (blue dashed). (c): Error measure, $E(\bar{f}_0)$, between the position of the leading edge given by the three-species heterogeneous model and the position predicted by the single-species homogeneous model as a function of amplitude of the interaction force, $\bar{f}_0$. (d)-(e): Cell density profiles predicted by the three-species heterogeneous model, $\mathcal{P}(x,t)$ (solid red), superimposed with density profiles given by the single-species homogeneous model calibrated with the best-fit value of $\bar{f}_0$, $P(x,t)$ (solid blue). The black arrow denotes the best-fit value of interaction force, $\bar{f}_0 = 0.06\,\mu\text{m/h}$. The continuum results for both models are presented at $t = 0, 12, 24, 36$, and $48\,\text{h}$. Black arrows denote the direction of increasing time. Results in (e) show a close-up comparison right near the leading edge, denoted by the gray shaded region in (d).

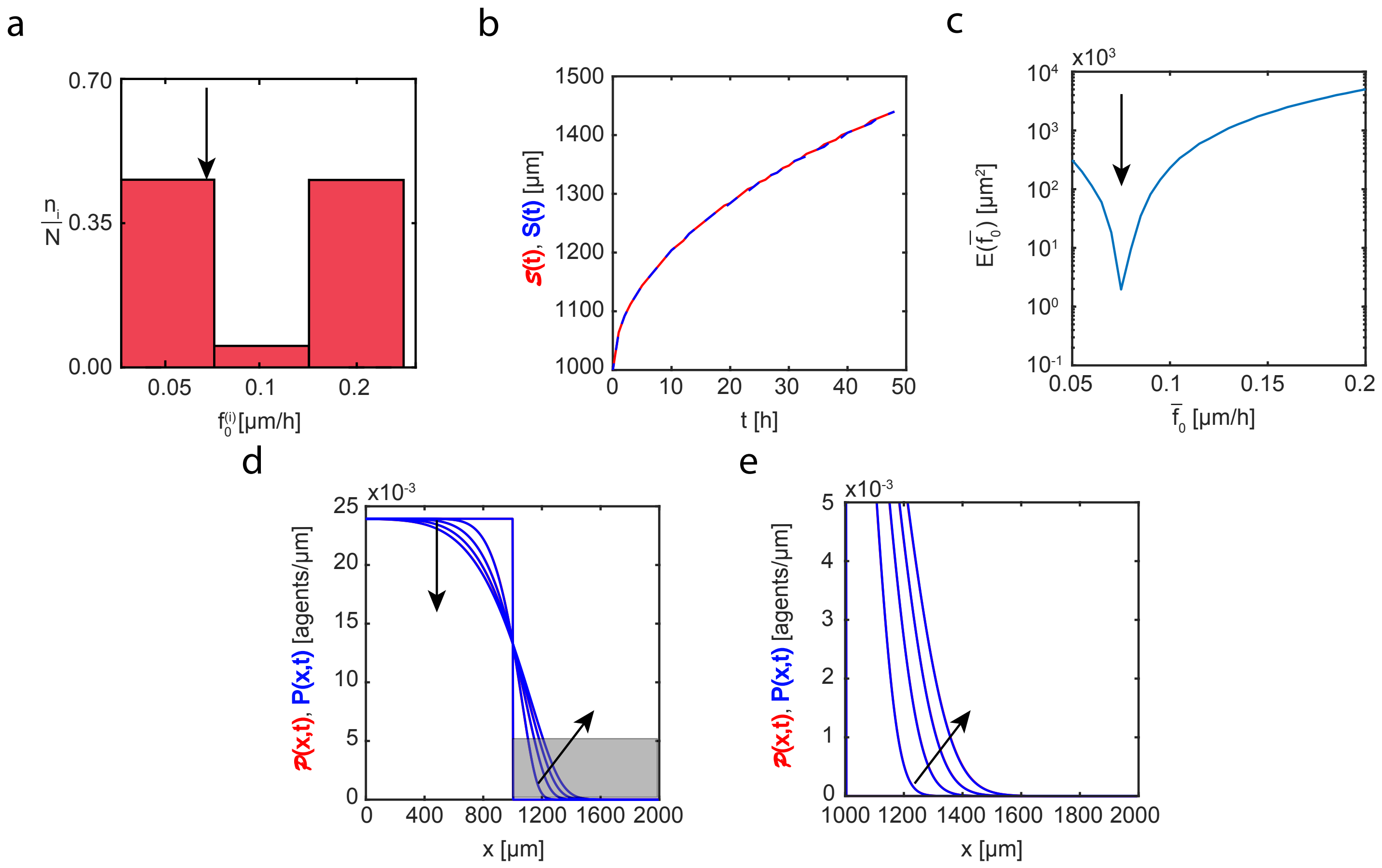


Fig. B.3. Set IIc. Heterogeneity in interaction forces: non-monotonic distribution. (a): Interaction force distribution adopted in the three-species heterogeneous model, Equations (3.2)-(3.4) (main document). Here the proportions of cells of different sizes are set to: (i) $n_1/N = 0.472$; (ii) $n_2/N = 0.056$; (iii) $n_3/N = 0.472$. (b): Leading edge predicted by the three-species heterogeneous model, $\mathcal{S}(t)$ (solid red), and the best-fit approximation given by the single-species homogeneous model, $S(t)$ (blue dashed). (c): Error measure, $E(\bar{f}_0)$, between the position of the leading edge given by the three-species heterogeneous model and the position predicted by the single-species homogeneous model as a function of amplitude of the interaction force, $\bar{f}_0$. (d)-(e): Cell density profiles predicted by the three-species heterogeneous model, $\mathcal{P}(x,t)$ (solid red), superimposed with density profiles given by the single-species homogeneous model calibrated with the best-fit value of $\bar{f}_0$, $P(x,t)$ (solid blue). The black arrow denotes the best-fit value of interaction force, $\bar{f}_0 = 0.065\,\mu\text{m/h}$. The continuum results for both models are presented at $t = 0, 12, 24, 36,$ and $48\,\text{h}$. Black arrows denote the direction of increasing time. Results in (e) show a close-up comparison right near the leading edge, denoted by the gray shaded region in (d).

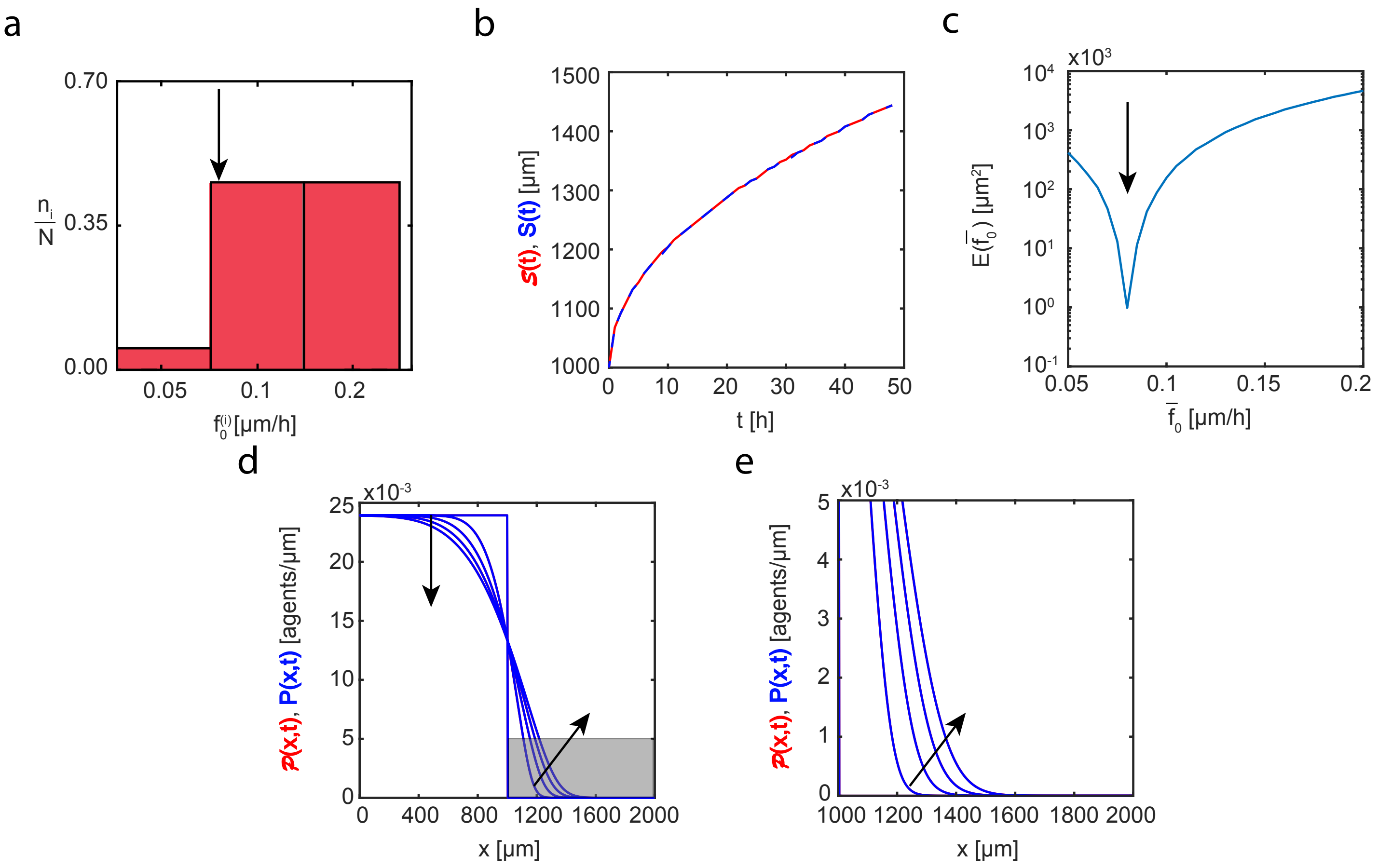


Fig. B.4. Set IId. Heterogeneity in interaction forces: monotonically increasing distribution. (a): Interaction force distribution adopted in the three-species heterogeneous model, Equations (3.2)-(3.4) (main document). Here the proportions of cells of different sizes are set to: (i) $n_1/N = 0.056$; (ii) $n_2/N = 0.472$; (iii) $n_3/N = 0.472$. (b): Leading edge predicted by the three-species heterogeneous model, $\mathcal{S}(t)$ (solid red), and the best-fit approximation given by the single-species homogeneous model, $S(t)$ (blue dashed). (c): Error measure, $E(\bar{f}_0)$, between the position of the leading edge given by the three-species heterogeneous model and the position predicted by the single-species homogeneous model as a function of amplitude of the interaction force, $\bar{f}_0$. (d)-(e): Cell density profiles predicted by the three-species heterogeneous model, $\mathcal{P}(x,t)$ (solid red), superimposed with density profiles given by the single-species homogeneous model calibrated with the best-fit value of $\bar{f}_0$, $P(x,t)$ (solid blue). The black arrow denotes the best-fit value of interaction force, $\bar{f}_0 = 0.07\,\mu$m/h. The continuum results for both models are presented at $t = 0, 12, 24, 36,$ and 48 h. Black arrows denote the direction of increasing time. Results in (e) show a close-up comparison right near the leading edge, denoted by the gray shaded region in (d).

### C Set III: Heterogeneity in the diffusivity

In this data set we explore the heterogeneity in the diffusivity where we fix values of the amplitude of the interaction force, $f_0^{(i)} = 0.05\,\mu$m/h, and the cell size, $\delta_i = 34\,\mu$m for $i = 1, 2, 3$. We note that these estimates are typical parameter values for PC-3 cells (Matsiaka et al., 2019). To analyse performance of the single-species homogeneous model (Equation (3.9), main document) applied to data generated by the three-species heterogeneous model (Equations (3.2)-(3.4), main document) we consider four different diffusivity distributions: (i) uniform distribution, Figure C.1(a), (ii) monotonically decreasing distribution, Figure C.2(a), (iii) non-monotonic distribution, Figure C.3(a), and (iv) monotonically increasing distribution, Figure C.4(a). For all cases presented we are able to predict a position of the leading edge as well as accurately describe cell density profiles given by the three-species heterogeneous model.

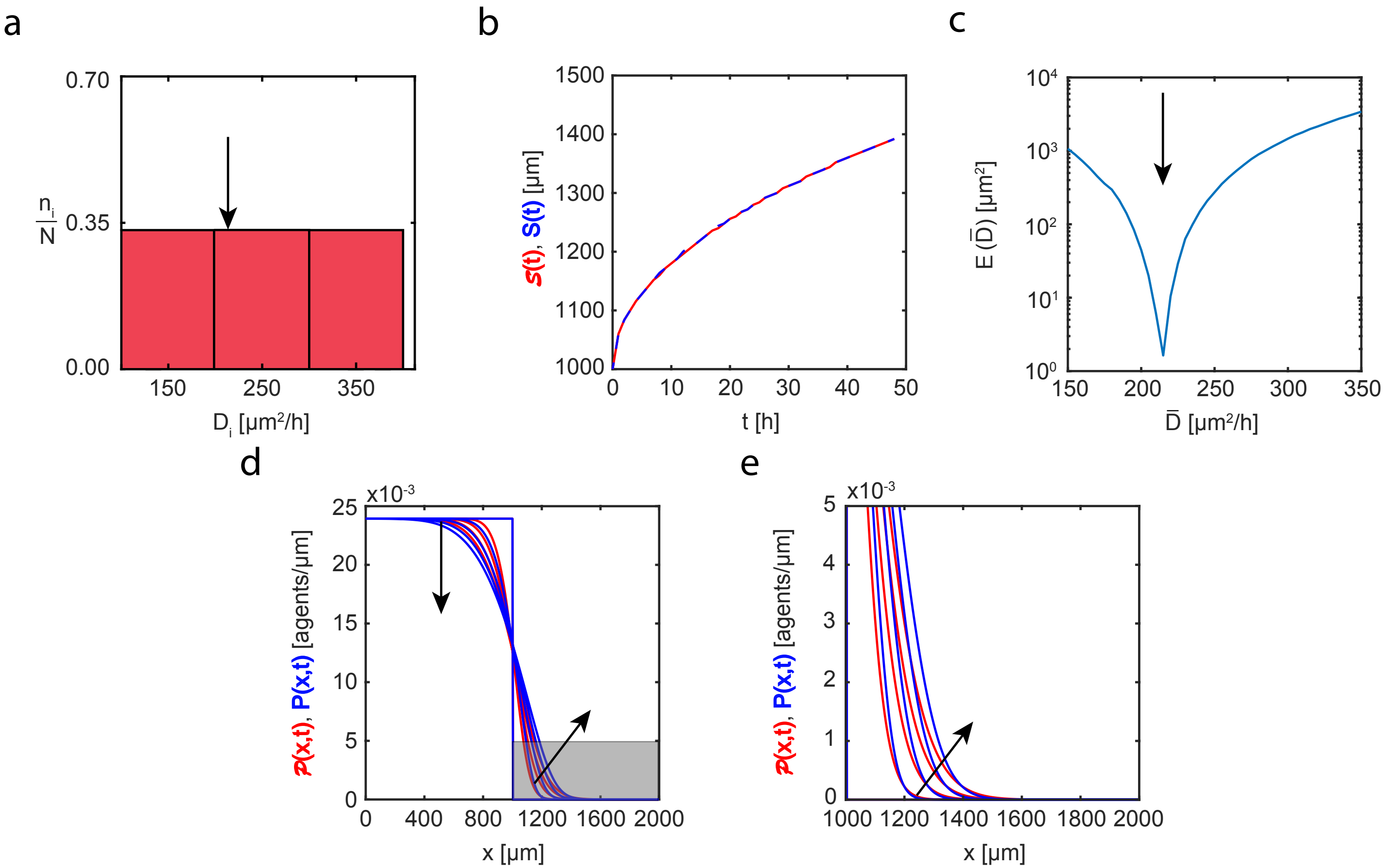


Fig. C.1. Set IIIa. Heterogeneity in diffusivity: uniform distribution. (a): Diffusivity distribution adopted in the three-species heterogeneous model, Equations (3.2)-(3.4) (main document). Here the proportions of cells of different sizes are set to: (i) $n_1/N = 0.333$; (ii) $n_2/N = 0.333$; (iii) $n_3/N = 0.333$. (b): Leading edge predicted by the three-species heterogeneous model, $\mathcal{S}(t)$ (solid red), and the best-fit approximation given by the single-species homogeneous model, $S(t)$ (blue dashed). (c): Error measure, $E(\bar{D})$, between the position of the leading edge given by the three-species heterogeneous model and the position predicted by the single-species homogeneous model as a function of diffusivity, $\bar{D}$. The black arrow denotes the best-fit value of diffusivity, $\bar{D} = 215\,\mu\text{m}^2/\text{h}$. (d)-(e): Cell density profiles predicted by the three-species heterogeneous model, $\mathcal{P}(x,t)$ (solid red), superimposed with density profiles given by the single-species homogeneous model calibrated with the best-fit value of $\bar{D}$, $P(x,t)$ (solid blue). The continuum results for both models are presented at $t = 0, 12, 24, 36,$ and $48\,\text{h}$. Black arrows denote the direction of increasing time. Results in (e) show a close-up comparison right near the leading edge, denoted by the gray shaded region in (d).

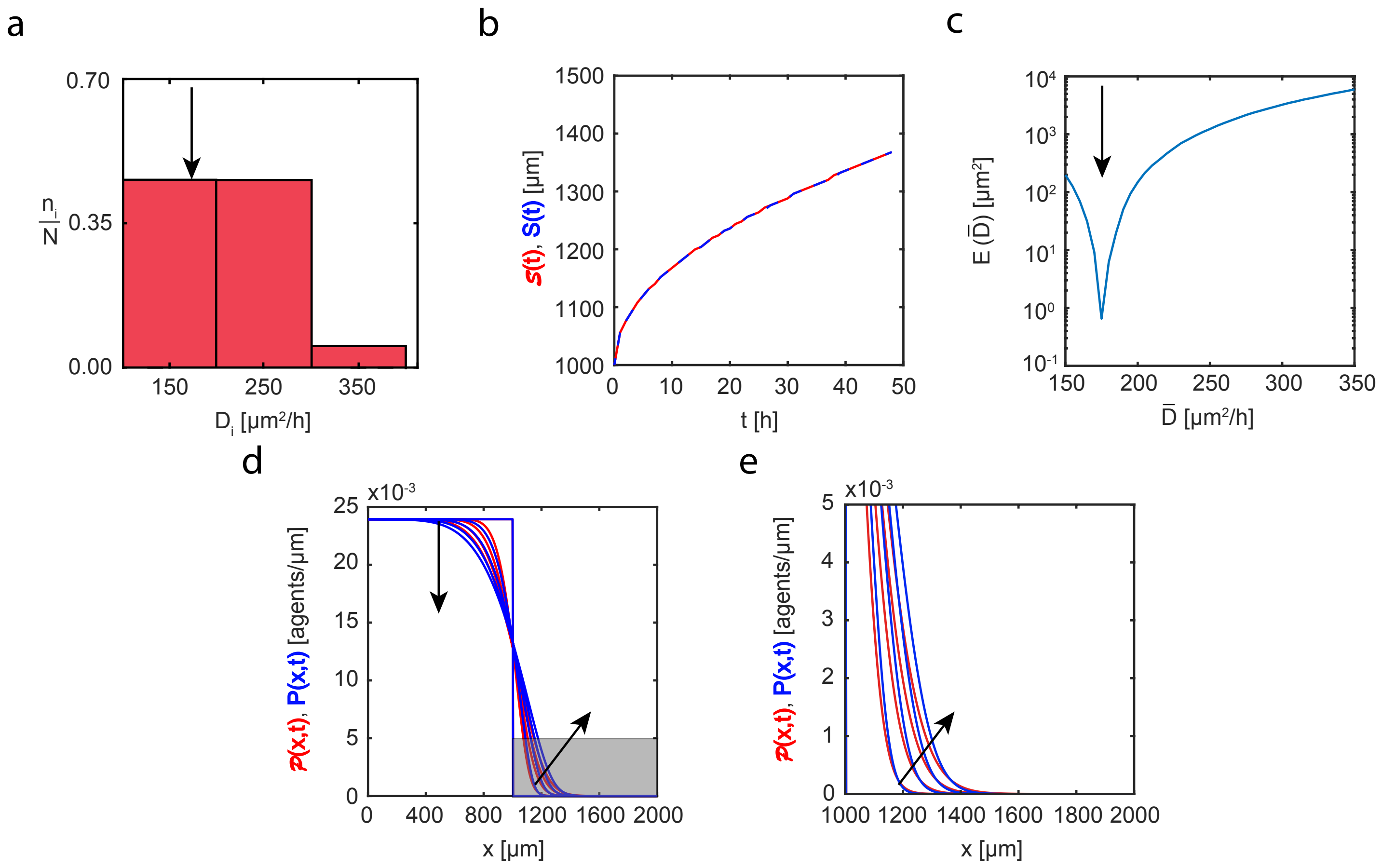


Fig. C.2. Set IIIb. Heterogeneity in diffusivity: monotonically decreasing distribution. (a): Diffusivity distribution adopted in the three-species heterogeneous model, Equations (3.2)-(3.4) (main document). Here the proportions of cells of different sizes are set to: (i) $n_1/N = 0.472$; (ii) $n_2/N = 0.472$; (iii) $n_3/N = 0.056$. (b): Leading edge predicted by the three-species heterogeneous model, $\mathcal{S}(t)$ (solid red), and the best-fit approximation given by the single-species homogeneous model, $S(t)$ (blue dashed). (c): Error measure, $E(\bar{D})$, between the position of the leading edge given by the three-species heterogeneous model and the position predicted by the single-species homogeneous model as a function of diffusivity, $\bar{D}$. The black arrow denotes the best-fit value of diffusivity, $\bar{D} = 175\,\mu\text{m}^2/\text{h}$. (d)-(e): Cell density profiles predicted by the three-species heterogeneous model, $\mathcal{P}(x,t)$ (solid red), superimposed with density profiles given by the single-species homogeneous model calibrated with the best-fit value of $\bar{D}$, $P(x,t)$ (solid blue). The continuum results for both models are presented at $t = 0, 12, 24, 36$, and $48\,\text{h}$. Black arrows denote the direction of increasing time. Results in (e) show a close-up comparison right near the leading edge, denoted by the gray shaded region in (d).

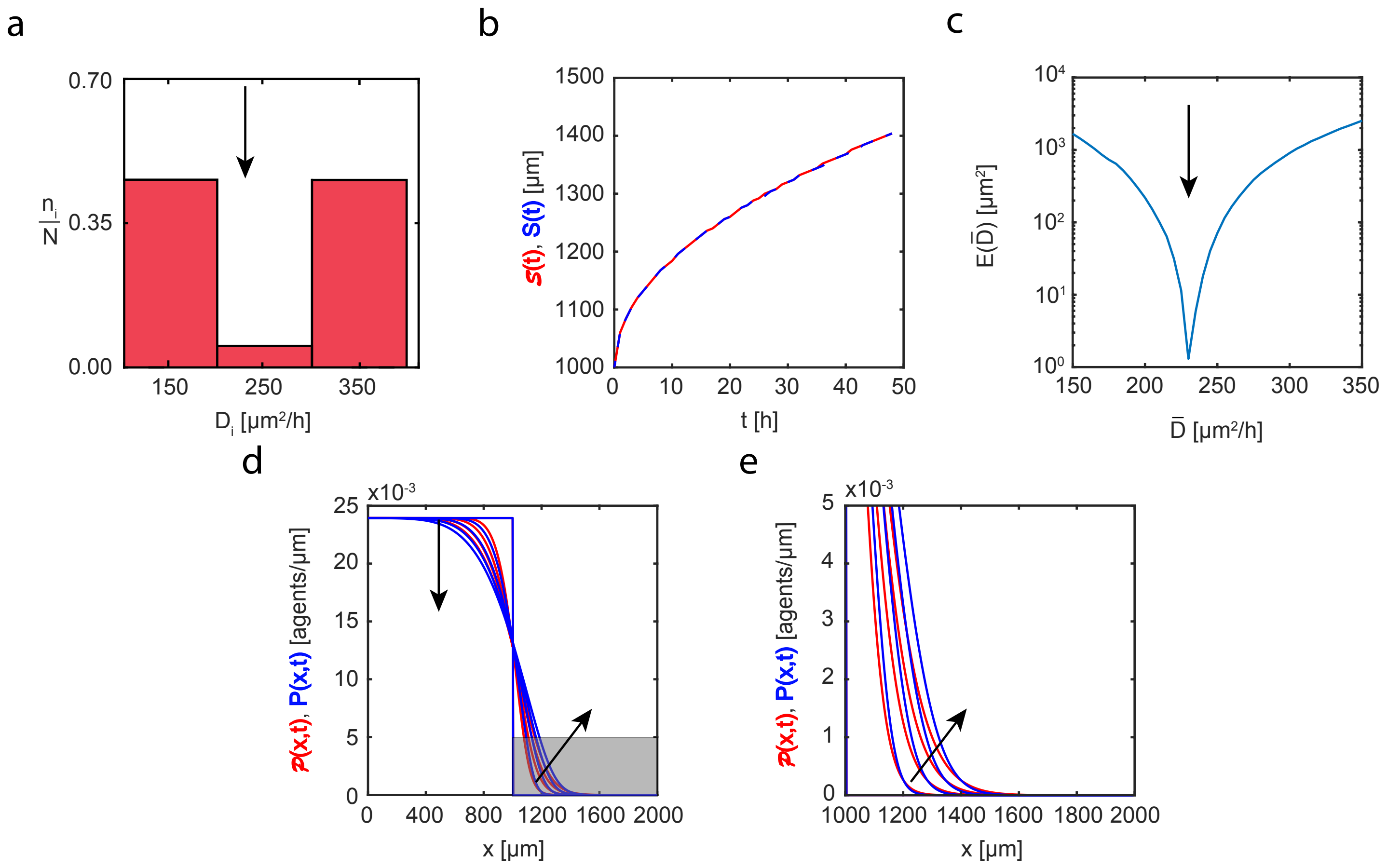


Fig. C.3. Set IIIc. Heterogeneity in diffusivity: non-monotonic distribution. (a): Diffusivity distribution adopted in the three-species heterogeneous model, Equations (3.2)-(3.4) (main document). Here the proportions of cells of different sizes are set to: (i) $n_1/N = 0.472$; (ii) $n_2/N = 0.056$; (iii) $n_3/N = 0.472$. (b): Leading edge predicted by the three-species heterogeneous model, $\mathcal{S}(t)$ (solid red), and the best-fit approximation given by the single-species homogeneous model, $S(t)$ (blue dashed). (c): Error measure, $E(\bar{D})$, between the position of the leading edge given by the three-species heterogeneous model and the position predicted by the single-species homogeneous model as a function of diffusivity, $\bar{D}$. The black arrow denotes the best-fit value of diffusivity, $\bar{D} = 230\,\mu\text{m}^2/\text{h}$. (d)-(e): Cell density profiles predicted by the three-species heterogeneous model, $\mathcal{P}(x,t)$ (solid red), superimposed with density profiles given by the single-species homogeneous model calibrated with the best-fit value of $\bar{D}$, $P(x,t)$ (solid blue). The continuum results for both models are presented at $t = 0, 12, 24, 36,$ and $48\,\text{h}$. Black arrows denote the direction of increasing time. Results in (e) show a close-up comparison right near the leading edge, denoted by the gray shaded region in (d).

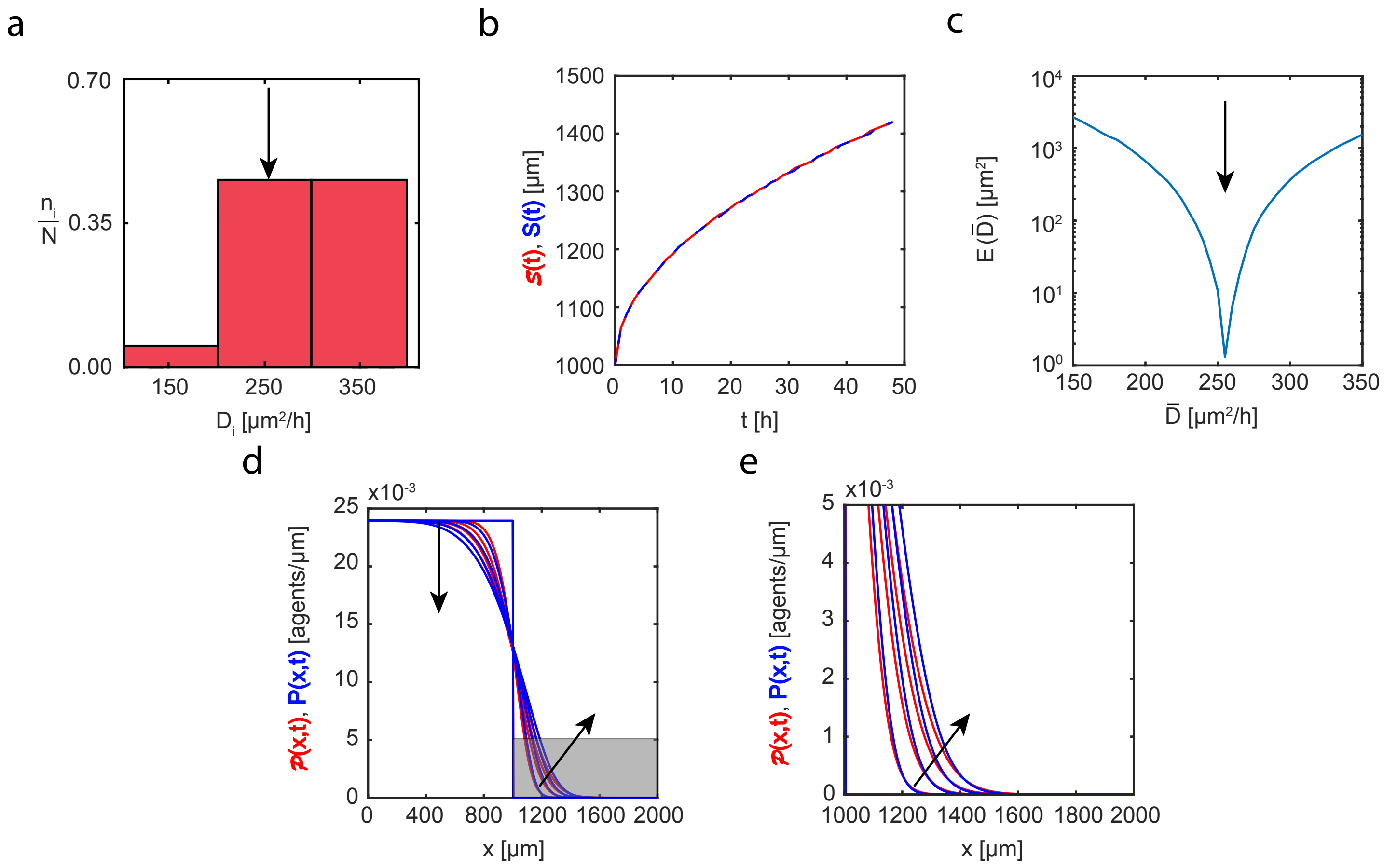


Fig. C.4. Set IIId. Heterogeneity in diffusivity: monotonically increasing distribution. (a): Diffusivity distribution adopted in the three-species heterogeneous model, Equations (3.2)-(3.4) (main document). Here the proportions of cells of different sizes are set to: (i) $n_1/N = 0.056$; (ii) $n_2/N = 0.472$; (iii) $n_3/N = 0.472$. (b): Leading edge as predicted by the three-species heterogeneous model, $\mathcal{S}(t)$ (solid red), and the best-fit approximation given by the single-species homogeneous model, $S(t)$ (blue dashed). (c): Error measure, $E(\bar{D})$, between the position of the leading edge given by the three-species heterogeneous model and the position predicted by the single-species homogeneous model as a function of diffusivity, $\bar{D}$. The black arrow denotes the best-fit value of diffusivity, $\bar{D} = 255\,\mu\text{m}^2/\text{h}$. (d)-(e): Cell density profiles as predicted by the three-species heterogeneous model, $\mathcal{P}(x,t)$ (solid red), superimposed with density profiles given by the single-species homogeneous model calibrated with the best-fit value of $\bar{D}$, $P(x,t)$ (solid blue). The continuum results for both models are presented at $t = 0, 12, 24, 36,$ and $48\,\text{h}$. Black arrows denote the direction of increasing time. Results in (e) show a close-up comparison right near the leading edge, denoted by the gray shaded region in (d).

### D Discretisation scheme for the single-species homogeneous model and heterogeneous three-species model

In this section we present the discretisation scheme used to obtain the numerical solution of the single-species homogeneous model in the mean-field framework. In summary, the governing equation that we consider is as follows,

$$\frac{\partial P(x,t)}{\partial t} = \bar{D}\Delta P(x,t) - (N-1)\nabla(P(x,t)\,V(x,t)), \tag{D.1}$$

where

$$V(x,t) = \int F(x-y)\,P(y,t)\,\mathrm{d}y \tag{D.2}$$

is the velocity filed induced by intercellular interaction forces, and $N = 36$ is the total number of cells in the simulations.

To present the numerical scheme as succinctly as possible, we define

$$\sigma(x,y,t) = F(x-y)\,P(y,t), \tag{D.3}$$

$$\begin{aligned} I_s &= P(x_s,t)\int \sigma(x_s,y)\,\mathrm{d}y \\ &= P(x_s,t)\frac{h}{2}\sum_i \Big[\sigma(x_s,y_{i+1}) + \sigma(x_s,y_i)\Big] + O(h^2), \end{aligned} \tag{D.4}$$

where the rectangle rule with step $h$ is used for numerical integration, and indices $s$ and $i$ denote the equally-spaced spatial mesh nodes.

Using the definitions in Equations (D.3)-(D.4), we apply the method of lines to Equation (D.1) and obtain the following system of coupled ordinary differential equations,

$$\frac{\mathrm{d}P_i}{\mathrm{d}t} = \frac{D}{h^2}\Big[P_{i+1} - 2P_i + P_{i-1}\Big] - (N-1)\frac{1}{2h}\Big[I_{i+1} - I_{i-1}\Big], \tag{D.5}$$

where index $i$ denotes a spatial mesh node. This systems of ordinary differential equations is solved using an explicit forward Euler algorithm with constant time steps of duration $\Delta t$.

Similarly, the three-species model is given by three coupled integro-PDEs in the following form,

$$\frac{\partial p^{(1)}}{\partial t} = D_1 \Delta p^{(1)} - (n_1 - 1)\nabla(p^{(1)}\, V^{(11)}) - n_2 \nabla(p^{(1)}\, V^{(12)}) - n_3 \nabla(p^{(1)}\, V^{(13)}), \tag{D.6}$$

$$\frac{\partial p^{(2)}}{\partial t} = D_2 \Delta p^{(2)} - (n_2 - 1)\nabla(p^{(2)}\, V^{(22)}) - n_1 \nabla(p^{(2)}\, V^{(21)}) - n_3 \nabla(p^{(2)}\, V^{(23)}), \tag{D.7}$$

$$\frac{\partial p^{(3)}}{\partial t} = D_3 \Delta p^{(3)} - (n_3 - 1)\nabla(p^{(3)}\, V^{(33)}) - n_1 \nabla(p^{(3)}\, V^{(31)}) - n_2 \nabla(p^{(3)}\, V^{(32)}), \tag{D.8}$$

$$V^{(lk)} = \int_\Omega F^{(lk)}(x-y)\, p^{(k)}(y,t)\, \mathrm{d}y. \tag{D.9}$$

We define

$$\sigma^{lk}(x,y,t) = F^{(lk)}(x-y)\, p^{(k)}(y,t), \tag{D.10}$$

$$\begin{aligned} I_s^{lk} &= p^{(l)}(x_s,t) \int \sigma^{lk}(x_s,y)\, \mathrm{d}y \\ &= p^{(l)}(x_s,t) \frac{h}{2} \sum_i \Big[\sigma^{lk}(x_s,y_{i+1}) + \sigma^{lk}(x_s,y_i)\Big] + O(h^2), \end{aligned} \tag{D.11}$$

where $k = 1,2,3$ is the subpopulation index, indices $i$ and $s$ denote the equally-spaced spatial mesh nodes, and $h$ is spatial discretisation step.

Using the definitions in Equations (D.10)-(D.11), we apply the method of lines to Equations (D.6)-(D.8) and obtain the following system of coupled ordinary differential equations,

$$\frac{\mathrm{d}p_i^{(k)}}{\mathrm{d}t} = \frac{D_k}{h^2}\Big[P_{i+1}^{(k)} - 2P_i^{(k)} + P_{i-1}^{(k)}\Big] - (n_k - 1)\frac{1}{2h}\Big[I_{i+1}^{kk} - I_{i-1}^{kk}\Big] - \sum_{l \neq k} n_l \frac{1}{2h}\Big[I_{i+1}^{lk} - I_{i-1}^{lk}\Big], \tag{D.12}$$

where $k = 1,2,3$ is the subpopulation index.
